# Developmental Correlates of Epigenetic and Polygenic Indices of Cognition and Educational Attainment from Birth to Young Adulthood

**DOI:** 10.64898/2026.04.01.715866

**Authors:** D. Fraemke, L. Paulus, I. Schuurmans, J.-H. Walter, D. Czamara, A. M. Schowe, A. deSteiguer, P. T. Tanksley, A. Okbay, B. Mönkediek, J. Instinske, M. M. Nöthen, C. K. L. Dißelkamp, A. J. Forstner, E. B. Binder, C. Kandler, F. M. Spinath, U. Lindenberger, M. Malanchini, C. A. M. Cecil, C. Mitchell, K. P. Harden, E. M. Tucker-Drob, L. Raffington

## Abstract

Large-scale genomic studies have identified biomarkers of adult cognitive functioning and educational attainment, yet the developmental pathways connecting these biomarkers to adult outcomes remain unclear. Drawing on four cohorts, we examined the developmental correlates of an epigenetic index of adult cognitive function (‘Epigenetic-*g*’) alongside polygenic indices of cognition and education. Epigenetic-*g* and polygenic indices were uncorrelated and captured distinct variation in children’s cognitive and academic performance. Longitudinal analyses revealed that Epigenetic-*g* is plastic in early childhood, reaching moderate stability by adolescence, and, unlike polygenic indices, is not related to longitudinal cognitive growth. Twin models indicated that Epigenetic-*g* captures genetic and unique environmental variation relevant to cognitive and academic achievement that is not identified by current polygenic indices. Epigenetic indices relevant to psychological development can be generated from DNA methylation studies of adults, with most variation in these indices emerging early in life.

## Main

Cognitive performance and educational attainment prospectively predict a range of socially consequential lifecourse outcomes, including socioeconomic status, dementia hazard, and life expectancy (Beam et al., 2020; Deary et al., 2010; Shenhar et al., 2026; Tucker-Drob, 2019). Large-scale studies of individual differences in adult cognitive function and educational attainment have discovered replicable genomic correlates, including both (genetic) DNA sequence variants and (epigenetic) DNA methylation sites (Lee et al., 2018; McCartney et al., 2022; Okbay et al., 2022). But adult cognitive function and educational attainment are the cumulative products of development, reflecting a decades-long interplay between genetic predispositions and the environments children encounter at home, in school, and society. Currently, the developmental pathways connecting genetic and epigenetic variation with cognitive function and educational attainment are poorly understood. Elucidating these pathways is essential to identifying periods of plasticity, selecting intervention targets, and determining appropriate uses of genomic and epigenomic biomarkers.

The developmental dynamics of DNA methylation differences known to be associated with adult cognitive function are especially poorly understood. One study of blood DNA methylation in approximately 9,000 Scottish adults, spanning ages 18 to 99 years old, found epigenetic variation that was significantly associated with a general factor of cognitive function (i.e., a “*g*” factor; McCartney et al., 2022). These discoveries were used to develop a biomarker, ‘Epigenetic-*g*’, which aggregates information about DNA methylation across the epigenome. In independent adult samples, including a racially diverse U.S. cohort, Epigenetic-*g* is correlated with cognitive performance (*R*^2^ ∼ 4%), brain structure, accelerated cognitive decline, and prospective dementia risk (Faul et al., 2025; McCartney et al., 2022). How Epigenetic-*g* changes across development, however, is largely unknown. In late childhood and adolescence, Epigenetic-*g* was found to have moderate rank-order stability across two measurement waves (deSteiguer et al., 2025). It is possible that Epigenetic-*g* becomes progressively more stable —*i.e.*, that it canalizes— as children mature into adolescents and young adults, as has been previously observed for cognitive function (Tucker-Drob et al., 2013; Tucker-Drob & Briley, 2014). Here, we investigate developmental change and stability in Epigenetic-*g* using longitudinal epigenetic data from four cohorts spanning birth to age 30.

A related question about Epigenetic-*g* is its relationship to genetic variation. As an epigenetic mechanism, DNA methylation regulates multi-level interactions between genes and their cellular and larger ecological environments in a highly age-dependent manner (Bertucci-Richter et al., 2024; Smith & Meissner, 2013). A DNA methylation biomarker could therefore largely reflect either genetic variation, environmental variation, or both. One previous study of children found that Epigenetic-*g* was moderately heritable (∼60%; Raffington et al., 2023). But it remains unknown to what extent Epigenetic-*g* is related to genetic variants previously linked to cognitive and academic performance. Genome-wide association studies have identified common genetic variants associated with cognitive function and educational attainment (Lee et al., 2018; Okbay et al., 2022). Polygenic indices (PGIs), which aggregate information about trait-associated variants across the genome, capture up to 10% of the variance in adult cognition, and up to 16% of the variance in educational attainment. They also predict children’s cognitive and academic performance (Allegrini et al., 2019; de Zeeuw et al., 2014; Selzam et al., 2017). If Epigenetic-*g* primarily reflects genetic variation, we expect positive correlations with education and cognition PGIs. If, on the other hand, it mainly captures highly plastic early-life environmental processes, we expect weak correlations with these PGIs. The extent to which epigenetic and genetic biomarkers are capturing distinct processes also bears on the question as to whether these biomarkers can predict unique variation in young people’s cognitive and academic performance.

Although PGIs are themselves immutable, their associations with academic achievement increase across adolescence (Harden et al., 2020; Malanchini et al., 2024). Whether a similar developmental dynamic is apparent for Epigenetic-*g* is unknown. A previous cross-sectional study in children and adolescents found that salivary Epigenetic-*g* was correlated with children’s performance on various cognitive and academic tasks, accounting for 11% of the variation in math performance (Raffington et al., 2023), but this association has not yet been investigated in longitudinal data. Given the potential malleability of DNA methylation across the lifespan, Epigenetic-*g* might be expected to be a more sensitive biomarker than DNA-based PGIs to change in cognition over time.

Finally, while epigenetic and genetic biomarkers aim to capture molecular variation, previous research on PGIs have found that they also capture socially stratified, environmental differences. When PGIs are applied within families, their associations with cognition and, especially, educational attainment are significantly attenuated, relative to associations observed in the general population. This suggests that many PGI associations are, at least in part, driven by environmentally mediated pathways correlated with genotype and varying between families (Demange et al., 2021; Selzam et al., 2019; Starr et al., 2025). For example, family socioeconomic status (SES) shapes developmental environments and correlates with genotype (Abdellaoui et al., 2025). Regarding Epigenetic-*g,* research has shown modest associations with family and neighborhood SES (Raffington et al., 2023), yet no study has tested whether Epigenetic-*g* effects show similar attenuation when applied within-family. Evidence of such attenuation would imply that shared family environments contribute to epigenetic variation, whereas persistence within families would point to direct genetic effects and sibling-specific environmental influences. Twin comparisons further sharpen this inference: Within-dizygotic-twin pair analyses control for family-level effects, while within-monozygotic-twin pair analyses test whether differences in Epigenetic-*g* covary with cognition when nuclear DNA, birth date, and shared background are held constant. The latter would indicate that Epigenetic-*g* is sensitive to idiosyncratic but meaningful environmental sources of variation.

Thus, this study addresses four major gaps in knowledge. *First*, it is unclear how Epigenetic-*g* changes over development. *Second*, the overlap between Epigenetic-*g* and PGIs for cognition and education remains unknown. *Third*, there is a lack of evidence regarding whether these biomarkers, particularly Epigenetic-*g*, are associated with longitudinal change in child cognitive development. And *fourth*, whether Epigenetic-*g* is associated with child cognitive and academic performance even when comparing within families remains unclear. To address these questions, we analyzed birth cohort, longitudinal, and twin data from four pediatric cohorts across three countries (N > 10,000). We examined Epigenetic-*g* from birth to young adulthood and compared it to PGI-Cognition and PGI-Education throughout. Because cohorts differed in age at methylation measurement and tissue type, analyses were conducted within cohort. **Figure 1** overviews cohorts and measures.

**Figure 1.**
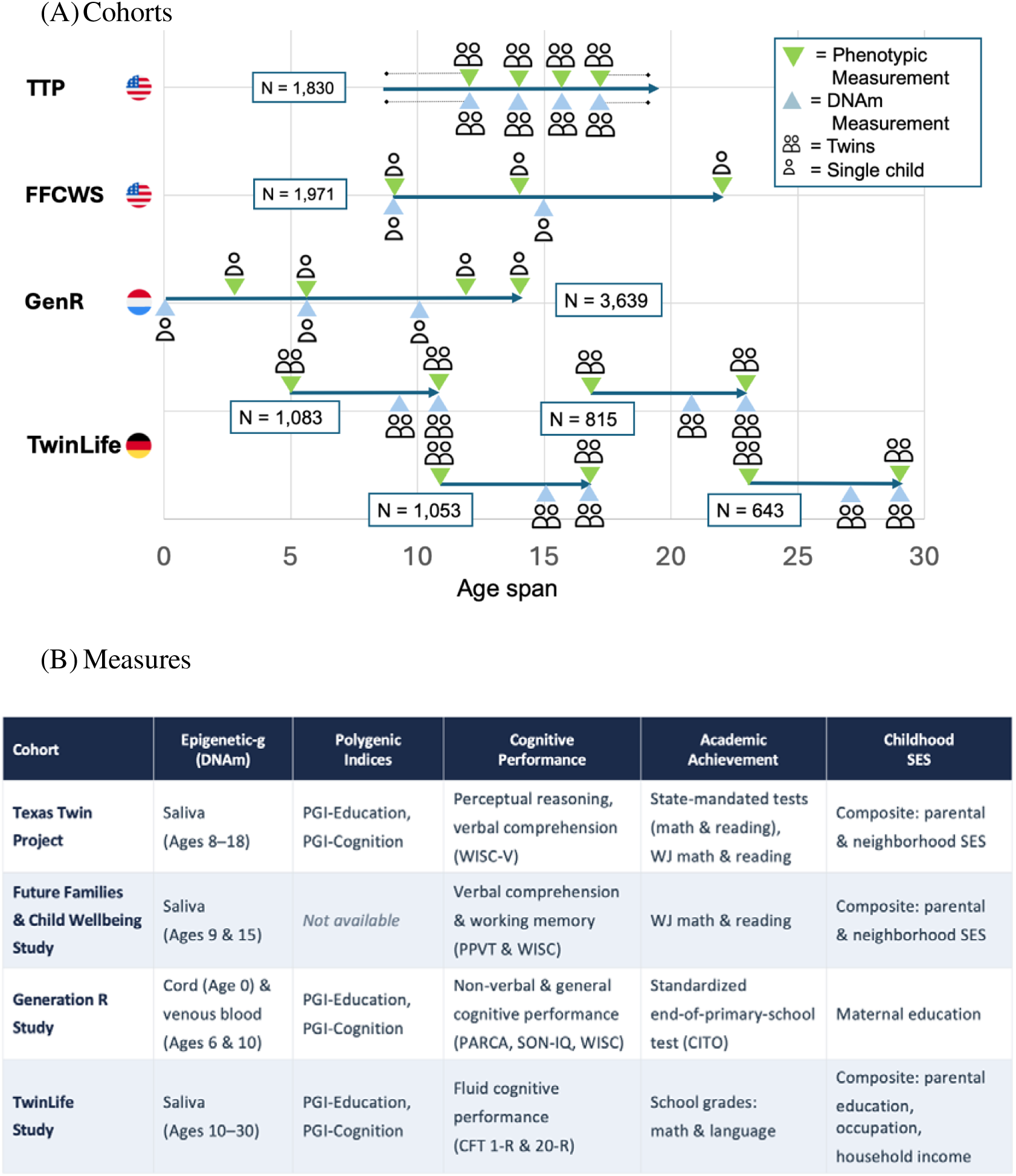
Overview of cohorts, measures, and analyses. **Panel A** displays the chronological age coverage of each cohort (Texas Twin Project [TTP], Future of Families and Child Wellbeing Study [FFCWS], The Generation R Study [GenR], and The TwinLife Study [TwinLife]). Horizontal lines indicate the observed age range and symbols mark assessment occasions; green inverted triangles denote phenotypic assessments and blue upright triangles denote DNA methylation (DNAm) collections, with icons indicating twin versus single-child samples and tissue source (saliva or blood). TwinLife consists of four sub-cohorts. Icons made via Freepik (www.flaticon.com). **Panel B** summarizes, by cohort, the availability of Epigenetic-*g* (including tissue type), polygenic indices, cognitive and academic outcome measures, and childhood socioeconomic status (SES) indicators. PGI = polygenic index; SES = socioeconomic status; PARCA = Parent Report of Children’s Abilities; CITO = Dutch standardized school achievement tests developed by Cito (Centraal Instituut voor Toetsontwikkeling); PPVT = Peabody Picture Vocabulary Test; WISC = Wechsler Intelligence Scale for Children; SON-IQ = Snijders-Oomen Non-verbal Intelligence Test; CFT 20-R = Culture Fair Test, 20-R.

**Figure 2.**
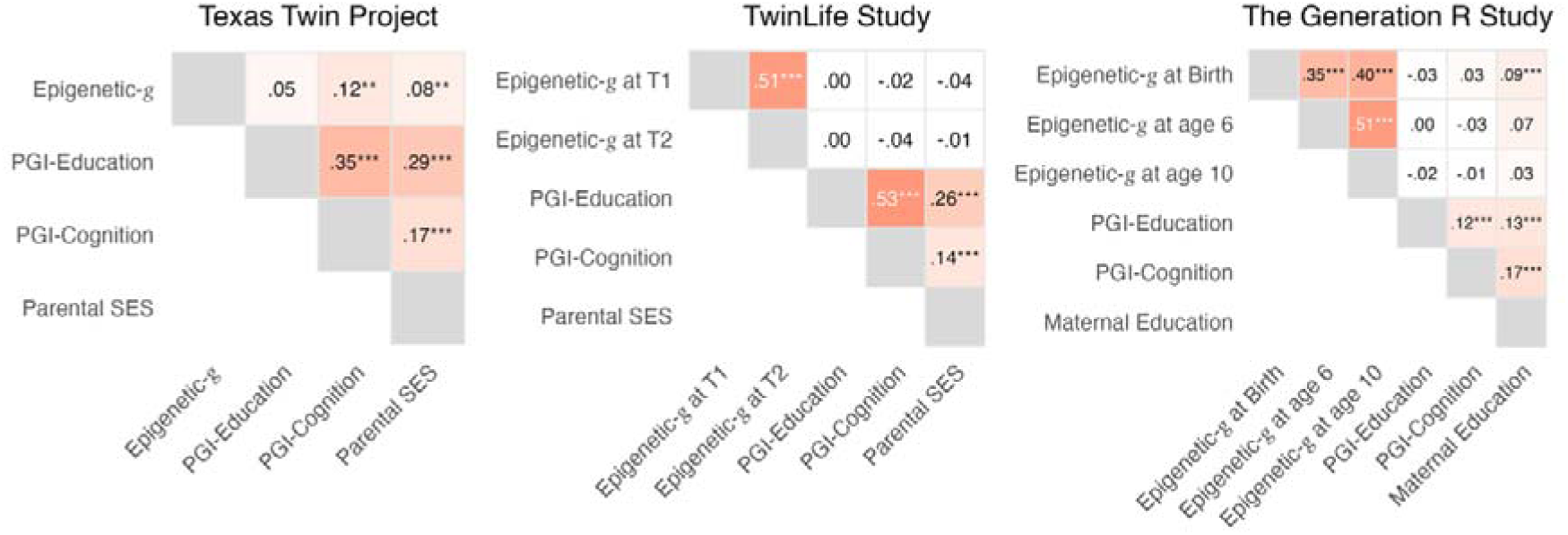
Associations between parental socioeconomic measures, epigenetic and polygenic indices of education and cognition in three developmental cohorts. Pairwise correlations among Epigenetic-*g* (at each available assessment wave), PGI-Education, PGI-Cognition, and parental socioeconomic indicators within the Texas Twin Project (TTP), TwinLife, and the Generation R Study (GenR). Correlations for the Future of Families and Child Wellbeing Study were not available. We report nominal *p* values: \**p* < .05. \*\**p* < .01. \*\*\**p* < .001.

## Results

### (1) How does Epigenetic-*g* Change over Development?

We examined how Epigenetic-*g* changes over development by testing whether it showed test-retest (rank order) stability across repeated measurements, whether its average level changed over time, and whether children differed in how rapidly they changed (Supplemental Table 1; see Supplemental Table 8 for descriptives of cohorts).

Regarding rank-order stability, repeated Epigenetic-*g* measures were moderately correlated in early and middle childhood, and more strongly correlated in later adolescence and young adulthood. In line with earlier findings from the Texas Twin Project (TTP; *r* = .34; deSteiguer et al., 2025), venous-blood levels at ages 6 and 10 were moderately correlated in Generation R (GenR) (*r* = .51, 95% CI [.43, .59], *p* < .001). Cord-blood Epigenetic-*g* at birth correlated moderately with Epigenetic-*g* at age 6 (*n* = 471; *r* = .36, 95% CI [.27, .43], *p* < .001) and age 10 (*n* = 440; *r* = .40, 95% CI [.32, .48], *p* < .001). In TwinLife, stability increased from moderate to strong across progressively older cohorts, with correlations between Epigenetic-*g* waves rising from cohort 1 (early adolescence; *n* = 360; *r* = .40, 95% CI [.31, .48]) to cohort 4 (young adulthood; *n* = 133; *r* = .62, 95% CI [.50, .72]), with intermediate estimates in cohort 2 (*n* = 393; *r* = .54, 95% CI [.46, .60]) and cohort 3 (*n* = 160; *r* = .57, 95% CI [.45, .66], all *ps* < .001).

Epigenetic-*g* tended to increase from childhood into adolescence (Supplemental Figures S1–S2). We modeled longitudinal methylation data using a latent difference score model in GenR (*n* = 327) and accelerated latent growth curve models in TTP (*n* = 1,327) and TwinLife (*n* = 1,055; Supplemental Table 1a). In GenR, the latent difference score model showed that Epigenetic-*g* increased on average from age 6 to 10 (mean change = 0.46 *SD* over four years, 95% CI [0.17, 0.75], *p* = .001). In TwinLife, the trend from age ∼10 to ∼30 was linear (unstandardized b = 1.15, 95% CI [0.89, 1.40], *p* < .001; Figure S2) while in TTP there was a positive quadratic age term (unstandardized b = 1.05, 95% CI [0.54, 1.55], *p* < .001; Figure S1). Across these different functional forms and age ranges, the parameters translate into an average increase of roughly 0.10 SD units of Epigenetic-*g* per year.

Moreover, we found that children differed in how rapidly Epigenetic-*g* changed when measured in blood across middle childhood, but not when measured in saliva across older children and adolescents. Specifically, we found significant variance in the latent difference between ages 6 and 10 in blood DNAm from GenR (σ²_ΔEpigenetic-*g*_ *=* 0.91, 95% CI [0.87, 0.94], *p* < .001). We did not find evidence for inter-individual differences in rates of change during adolescence; the variance of the Epigenetic-*g* slope factor was not significantly different from zero in TTP (σ²_slope_ = 0.06, 95% CI [–0.14, 0.26], *p* = .550) and TwinLife (σ²_slope_ = 0.07, 95% CI [–0.17, 0.32], *p* = .559; Supplemental Table 1a).

### (2) How are Epigenetic-*g* and PGIs Correlated Cross-Sectionally with Cognition, Education, and with Each Other?

The second set of analyses examined the unique and joint associations of Epigenetic-*g* and PGIs with cognitive and academic performance and how they are correlated with each other. As shown in Figure 3, Panel A, Epigenetic-*g* was associated with performance on at least one non-verbal cognitive task across all four cohorts, after FDR correction for multiple comparisons. In TTP and FFCWS there were also associations with higher performance on other cognitive measures (β range from 0.12 to 0.35, *p*_FDR_ ≤ .03; Figure 3; Supplemental Table 2). Moreover, Epigenetic-*g* was associated with all academic achievement measures in the TTP (β range from 0.18 to 0.34, *p*_FDR_ ≤ .007) and with WJ Reading in FFCWS (β = 0.42, *p*_FDR_ = .01), whereas the association with WJ Math in FFCWS was weaker and did not survive FDR correction (β = 0.23, *p*_FDR_ = .06). A follow-up analysis in TTP indicated that these associations extend to students’ educational pathways. Children enrolled in optional advanced math courses showed higher Epigenetic-*g* levels than those in basic courses (Welch’s *t*(46.75) = –5.22, *p* < .001, Cohen’s *d* = –0.74, 95% CI [–0.96, –0.42]; Figure 3, Panel B; Supplemental Table 2c).

**Figure 3.**
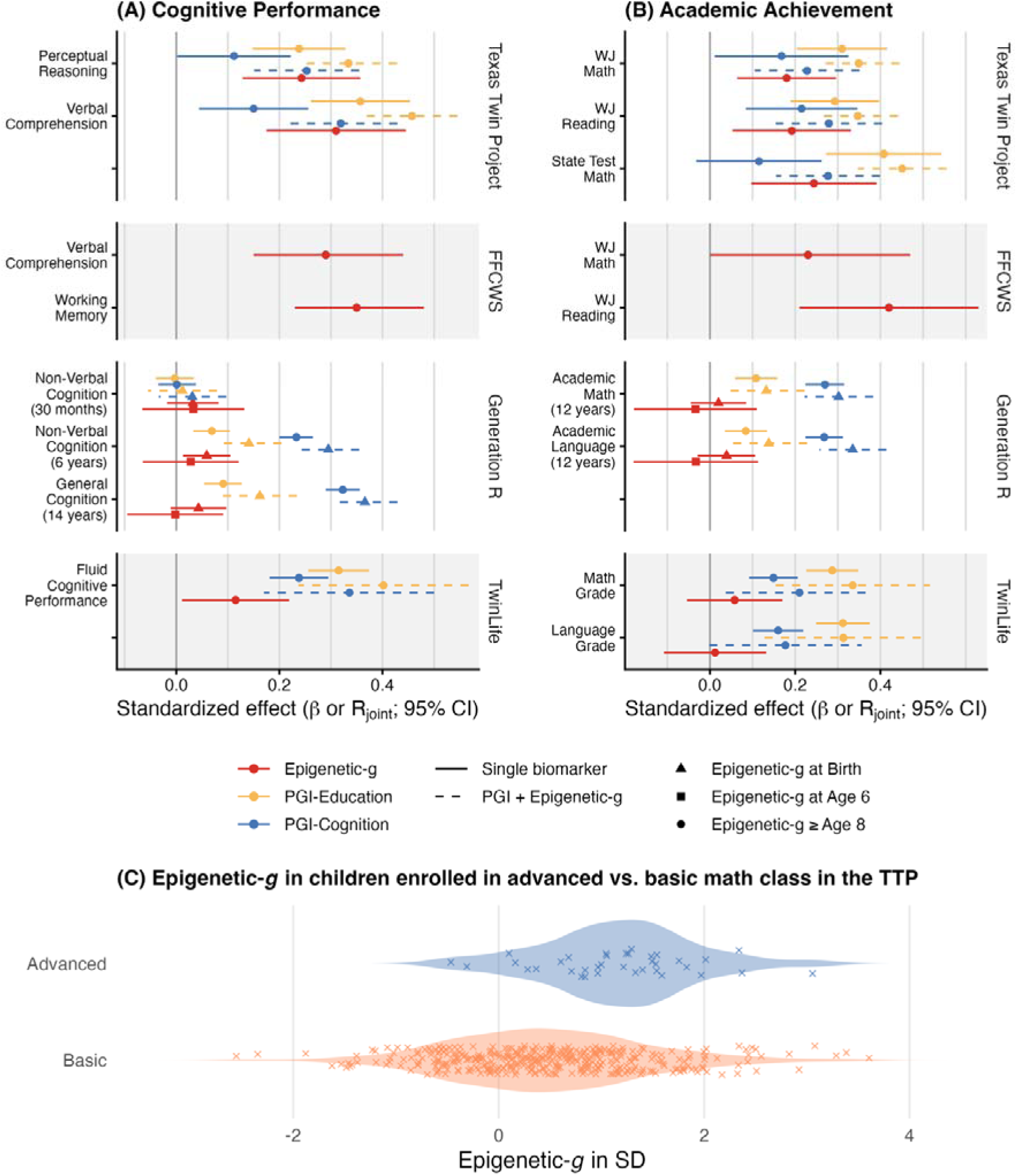
Cross-sectional associations of epigenetic and polygenic indices with cognitive performance in **Panel A** and academic achievements in **Panel B** across four cohorts. Panel A and B show standardized regression coefficients (β) for individual signals and partial multiple correlation coefficients (R_joint_) for the combination of two biomarkers. Estimates are derived from differing sample sizes as reported in descriptives in the supplemental material. Error bars indicate 95% confidence intervals. **Panel C** displays the distribution of Epigenetic-*g* by math class placement (advanced vs. basic) in the Texas Twin Project; points represent individual observations.

In the younger and smaller GenR cohort, evidence was somewhat less consistent: Associations between blood Epigenetic-*g* at age 6 with cognitive measures were in the expected direction, but not statistically significant (*p*_FDR_ ≥ .18). Notably, in the larger cord-blood GenR sample, Epigenetic-*g* at birth was modestly associated with non-verbal cognition at age 6 (β = 0.06, 95% CI [0.01, 0.11], *p*_FDR_ = .03). Neither Epigenetic-*g* at birth nor at age 6 was significantly related to academic achievement at age 12 (β range from −0.03 to 0.04, *p*_FDR_ ≥ .52).

Across all cohorts with PGIs (TTP, TwinLife and GenR), higher PGI-Education was associated with higher cognitive (β range from 0.07 to 0.36, *p*_FDR_ < .001) and academic performance (β range from 0.08 to 0.41, *p*_FDR_ < .001). Consistent with previous studies, PGI-Cognition showed a similar pattern in TTP and TwinLife, although with roughly half the effect sizes compared to cognitive outcomes (β range from 0.11 to 0.24, *p*_FDR_ < .05) and less consistent associations with academic performance (β range from 0.12 to 0.22, *p*_FDR_ range from < .001 to .13). In GenR, PGI-Cognition was more strongly related to most cognitive and academic measures than PGI-Education (β range from 0.23 to 0.32, *p*_FDR_ < .001), except for the earliest non-verbal cognitive test at 30 months, when neither PGI showed meaningful associations (*p*_FD_R ≥ .88).

In joint models, each PGI was included simultaneously with Epigenetic-*g* to assess their combined associations (Figure 3, Panel A and B; Supplemental Table 2 a–b). We report the partial multiple correlation, R_joint_, which quantifies the variance in the outcome jointly explained by Epigenetic-*g* and the corresponding PGI. When Epigenetic-*g* was associated with an outcome in the single-biomarker models, the corresponding joint models showed a stronger combined association than either biomarker alone, ranging from R_joint_ = 0.14 to 0.46 (*p*_FDR_ ≤ .04; Supplemental Table 2a–2b). Thus, Epigenetic-*g* and PGIs account for predominantly independent variation in cognitive and academic performance.

Additionally, associations between Epigenetic*-g* and PGIs were null to small (Supplemental Table 2d–e). PGI-Education was not significantly associated with Epigenetic-*g* in any cohort (β = –0.04 to 0.05, all *p* ≥ .284; PGIs unavailable in FFCWS). PGI-Cognition was modestly correlated with Epigenetic-*g* only in TTP (β = 0.13, 95% CI [0.003, 0.257], *p* = .045) and not significantly correlated in TwinLife and GenR (both β*s* < 0.01, *ps* ≥ .84).

### (3) Are Epigenetic-*g* and PGIs Associated with Longitudinal Change in Cognition and Academic Achievement?

Our third set of analyses examined associations with longitudinal change in cognitive and academic performance in TTP and TwinLife (Figure 4; Supplemental Table 3). Latent growth curve models decomposed longitudinal data for each outcome into an intercept (each child’s baseline level) and a slope (rate of within-person change), which we regressed on biomarkers.

**Figure 4.**
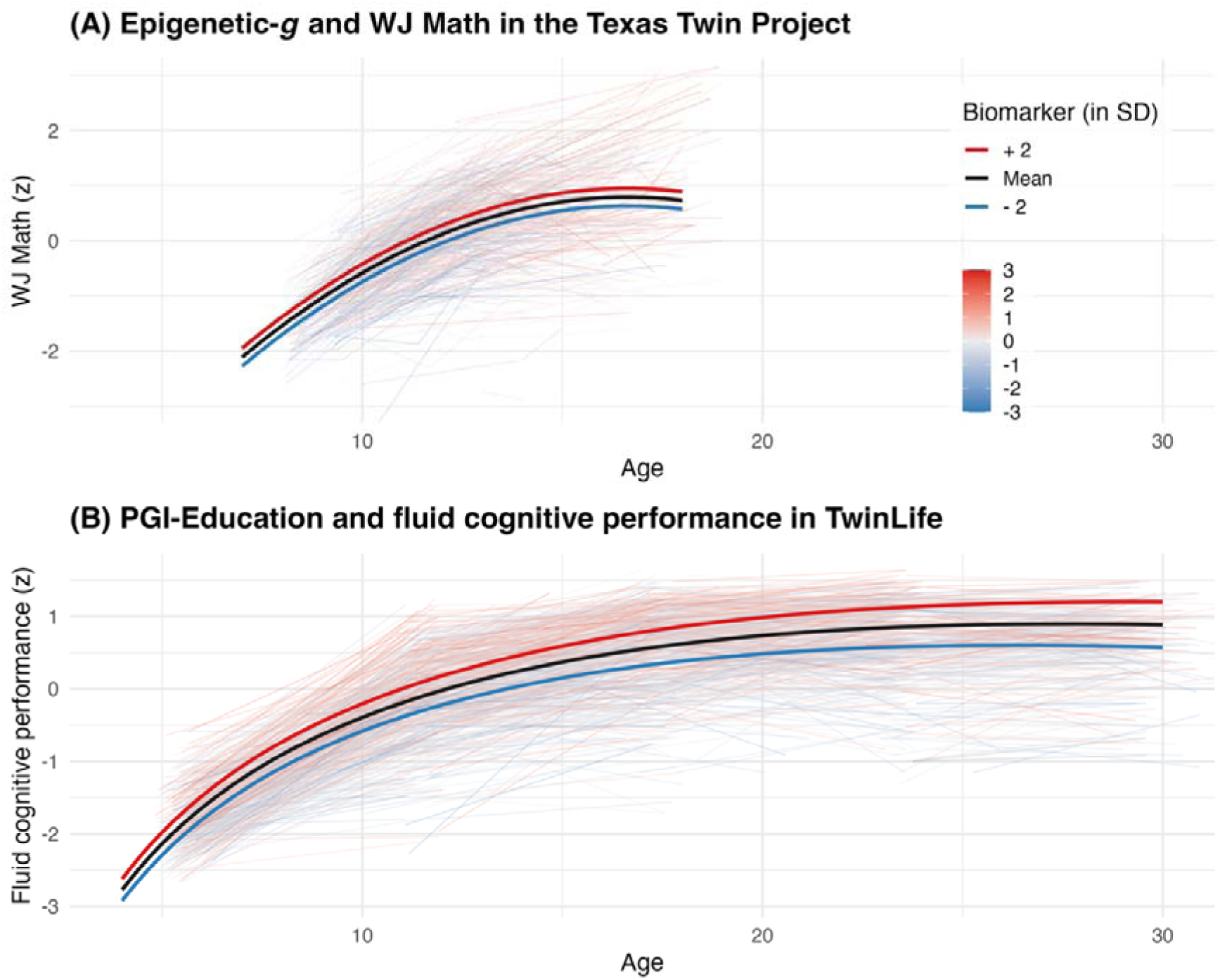
Selected longitudinal associations between polygenic and epigenetic indices with cognitive development. Panel A shows WJ Math over age in the Texas Twin Project (TTP). Thin lines depict individual trajectories (colored by baseline Epigenetic-*g*), and thick lines show model-implied trajectories at −2 SD, mean, and +2 SD of baseline Epigenetic-*g*. Panel B shows fluid cognitive performance over age in TwinLife, with individual trajectories colored by PGI-Education and model-implied trajectories plotted at −2 SD, mean, and +2 SD of PGI-Education. Panels A and B depict Epigenetic-*g* results in TTP and PGI results in TwinLife, respectively, because each cohort provides the largest available sample with the relevant repeated assessments, supporting comparatively stable visualization of developmental associations (or lack thereof). Full results, including models not visualized here, are reported in Supplemental Table 3. PGI = polygenic index; SD = standard deviation.

Across cohorts, baseline Epigenetic-*g* was not associated with developmental change in any outcome (*p*_FDR_ ≥ .49), suggesting that its association with cognitive and academic performance was stable across the observed developmental periods (Figure 4, Panel A).

In contrast, PGI-Education and PGI-Cognition were positively associated with higher average levels as well as inter-individual differences in intra-individual change in fluid cognitive performance in *n* = 3,008 children from TwinLife (PGI-Education on slopes: β = 0.44, 95% CI [0.15, 0.72], *p* = .003; PGI-Cognition: β = 0.42, 95% CI [0.15, 0.69], *p* = .002). In contrast, neither PGI was associated with developmental change in academic performance in TwinLife (*p*_FDR_ ≥ 0.26). In the smaller TTP cohort of *n* = 661 children, PGI-Education was associated with change in WJ Math (β = 0.22, 95% CI [0.05, 0.39], *p*_FDR_ = .02), but not with change in State Test Math, State Test Reading, WJ Reading or cognitive performance (*p*_FDR_ ≥ .59). PGI-Cognition did not show evidence of association with developmental change in any cognitive or academic outcome in the TTP (*p*_FDR_ ≥ .29).

Given polygenic associations with longitudinal change over time, we examined whether PGIs were also associated with school-track placement (TwinLife) and home learning indicators (TTP), which may reflect active and evocative gene-environment correlations in academic development over time (Figure 5; Supplemental Table 3c). In TwinLife, children with higher PGIs were more likely to be enrolled in the highest school track in Germany (PGI-Education: Welch’s *F*(2, 67.31) = 45.07, *p* < .001, η*²* = .05; PGI-Cognition: Welch’s *F*(2, 67.60) = 20.97, *p* < .001, η*²* = .02). In TTP, children with higher PGI-Education had greater home learning resources (β = 0.27, *p*_FDR_ < .001), fewer self-reported difficulties in math (β = −0.15, *p*_FDR_ = .013) and reading (β = −0.21, *p*_FDR_ = .002), and more reading outside school (β = 0.21, *p*_FDR_ < .001). PGI-Cognition showed a more selective pattern, with evidence primarily for fewer reported reading difficulties (β = −0.24, *p*_FDR_ = .004), and an association with reading outside school, which did not survive FDR correction (β = 0.18, *p*_FDR_ = .056).

**Figure 5.**
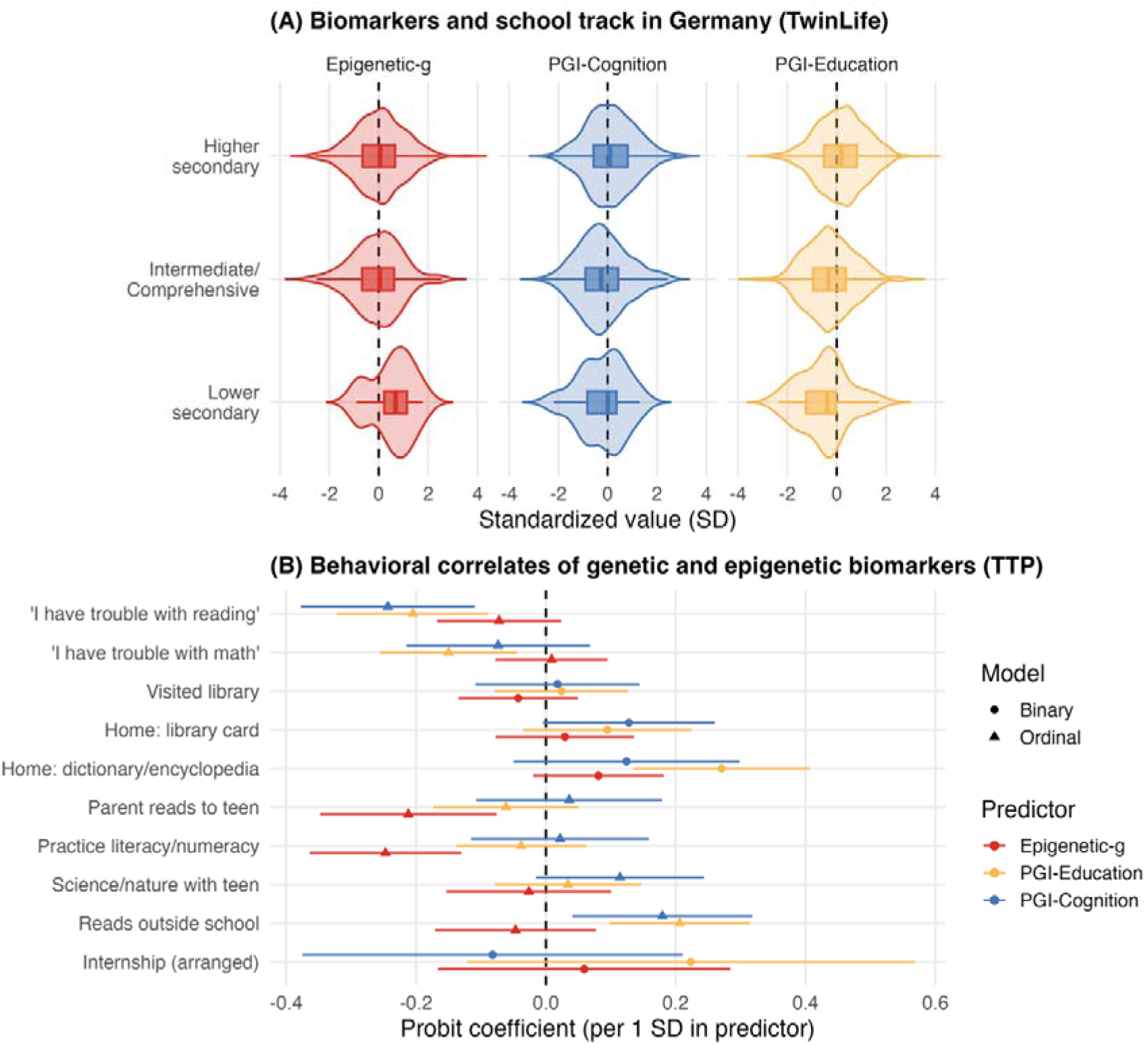
Cross-sectional associations of epigenetic and genetic biomarkers with school track and home learning. **Panel A** displays distributions of standardized biomarker values by German secondary school track in TwinLife (lower secondary, intermediate/comprehensive, higher secondary); these tracks broadly differ in academic orientation and typical qualifications, with lower secondary generally oriented toward vocational pathways and higher secondary (Gymnasium; roughly comparable to grammar-school) typically leading to the Abitur (university entrance qualification). Violins show the full distribution and embedded boxplots summarize central tendency and dispersion. **Panel B** presents associations of genetic and epigenetic biomarkers with adolescent academic involvement indicators in TTP as probit/ordered-probit coefficients per 1 *SD* increase in the predictor with 95% CIs (error bars); circles indicate binary outcomes and triangles indicate ordinal outcomes, with the dashed vertical line marking zero association. Full results are reported in Supplemental Table 3. *SD* = standard deviation; TTP = Texas Twin Project.

In contrast, Epigenetic-*g* did not vary by school track in TwinLife, Welch’s *F*(2, 21.81) = 2.19, *p* = .135, η*²* = .006. In TTP, higher Epigenetic-*g* was associated with less frequent parent-reported reading to the adolescent (β = −0.21, *p*_FDR_ = .011) and less frequent literacy/numeracy practice (β = −0.25, *p*_FDR_ < .001), whereas other associations were small and not significantly different from zero.

### (4) What are the Genetic and Environmental Contributions to Biomarker Associations?

Our fourth set of analyses probed the genetic, shared, and unique environmental contributions to biomarker associations. Specifically, we examined if associations persist within families, and to what extent they are stratified by childhood socioeconomic context. We tested for within-family attenuation using dizygotic twin pairs from TTP and TwinLife (Supplemental Table 4a – 4b). In TTP, Figure 6 shows that Epigenetic-*g* associations declined by ∼50% in within-family analyses. Yet, despite wider confidence intervals in the models, within-family associations remained statistically different from zero for two cognitive outcomes (perceptual reasoning: β = 0.14, 95% CI [0.04, 0.24], *p* = .005; verbal comprehension: β = 0.17, 95% CI [0.07, 0.27], *p* = .001) and WJ Math (β = 0.12, 95% CI [0.00, 0.24], *p* = .043). In TwinLife, between-family Epigenetic-*g* associations were small, and within-family estimates were of a similar magnitude, providing little evidence of within-family attenuation. Consistent with prior polygenic studies, effect sizes of PGI-Education predicting cognition were attenuated by roughly 50% in dizygotic twin analyses, whereas attenuation was less pronounced for PGI-Cognition.

**Figure 6.**
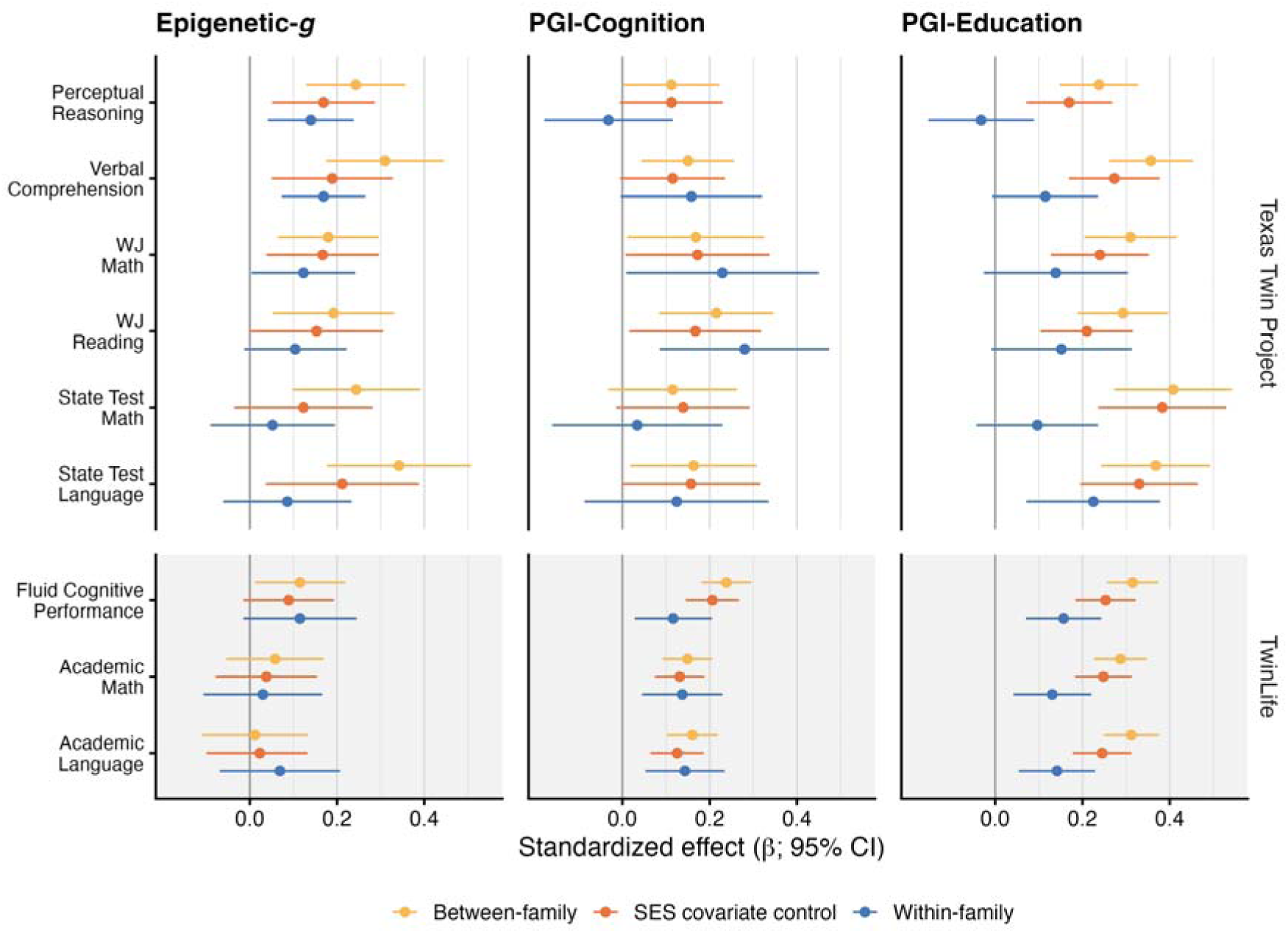
Within- and between-family analyses of epigenetic and polygenic associations with cognitive and academic outcomes in the Texas Twin Project and TwinLife. Standardized associations (β) are shown for Epigenetic-*g*, PGI-Cognition, and PGI-Education across cognitive and academic measures. Estimates are presented for between-family models (yellow), between-family models additionally adjusting for childhood socioeconomic status (SES; orange), and within-family dizygotic twin models (blue). Points indicate standardized effect estimates and horizontal lines indicate 95% confidence intervals; the vertical reference line marks zero association. Cohort labels on the right denote the Texas Twin Project (upper panels) and TwinLife (lower panels). PGI = polygenic index; SES = socioeconomic status.

Next, we computed twin models comparing the relative similarity of dizygotic and monozygotic twin pairs in Epigenetic-*g* (Supplemental Table 4c; Neale & Maes, 2004). Epigenetic-*g* showed moderate heritability in both cohorts (TTP: *h*² = .588, *p* < .001, *n* = 347 MZ and 656 DZ pairs; TwinLife: *h*² = .613, *p* < .001, *n* = 611 MZ and 713 DZ pairs), with the remaining variance attributable to unique environmental influences not shared by siblings, including measurement error (TTP: *e*² = .412, *p* < .001; TwinLife: *e*² = .387, *p* < .001).

We then tested whether genetic and unique environmental influences accounted for associations between Epigenetic-*g* and cognition (*i.e*., Cholesky twin model decompositions; Supplemental Tables 4d–4e). In *N* = 1,003 twin pairs from TTP, there was a genetic correlation between Epigenetic-*g* and Verbal Comprehension (*rA* = .573, *p* = .002). In TwinLife (*n* = 1,324 pairs), there was no significant genetic correlation with any cognitive or academic measure (*ps* > .09). Moreover, there were weak unique environmental associations between Epigenetic-*g* and perceptual reasoning, State Test Math, State Test Reading and WJ Math in TTP, as well as fluid cognition in TwinLife, although estimates were not significantly different from zero (*r*Es range from 0.115 to 0.170; *ps* range from .054 to .119).

As a complementary analysis, we examined monozygotic (MZ) twin-difference associations between Epigenetic-*g* and cognition (Figure 7; Supplemental Tables 4f–4g). In TTP, MZ differences in Epigenetic-*g* correlated with MZ differences in abilities that were more related to the fluid domain, such as perceptual reasoning (β = 0.128, 95% CI [0.01, 0.25], *p* = .038; *n* = 221 pairs), WJ Math (β = 0.177, 95% CI [0.02, 0.34], *p* = .029; *n* = 147 pairs), and State Test Math (β = 0.181, 95% CI [0.01, 0.35], *p* = .040; *n* = 120 pairs). In contrast, correlations with verbal skills were not significantly different from zero (verbal comprehension: *p* = .104, *n =* 222 pairs; WJ Reading: *p* = .673; *n* = 148 pairs, State Test Reading: *p* = .328, *n =* 120 pairs). In TwinLife, no MZ twin-difference association was statistically significant (*p* ≥ .083; *n* pairs range from 188 to 243).

**Figure 7.**
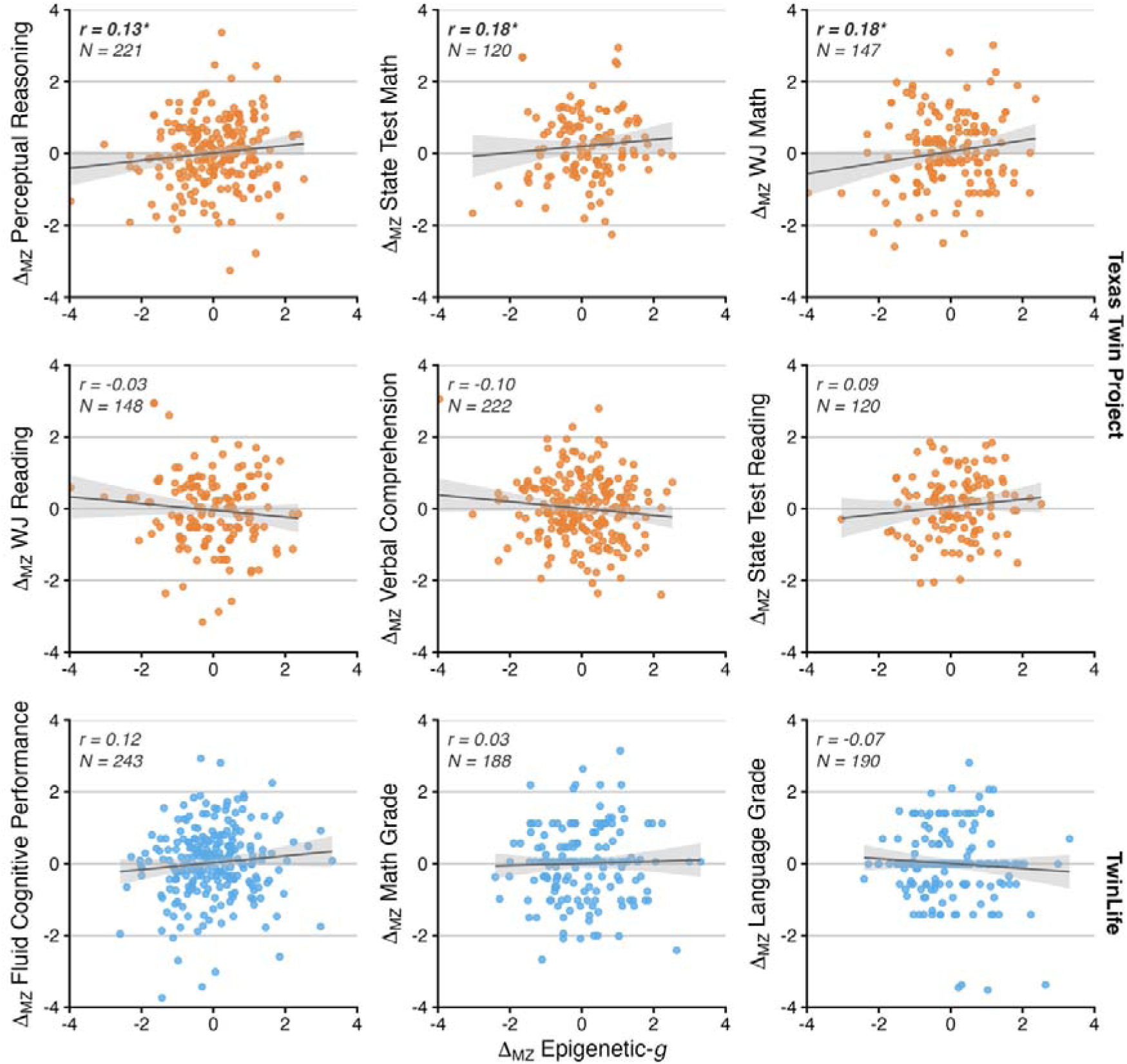
Associations of monozygotic twin differences () in Epigenetic-*g* and cognitive or academic outcomes. Each panel depicts the association between within-pair differences in Epigenetic-*g* (in SD) and within-pair differences in performance (in SD), thereby isolating covariation independent of additive genetic differences and shared family influences. Points represent standardized MZ twin-pair differences; solid lines show fitted linear regression estimates with 95% confidence bands. Pearson correlations (*r*) and the number of MZ pairs (*N*) are reported within panels; asterisks denote nominal significance (*p* < .05). Upper two orange panels correspond to the Texas Twin Project and the lower blue panel corresponds to TwinLife.

Next, we examined whether biomarkers were correlated with childhood socioeconomic status (SES; Figure 2) and whether biomarker-outcome associations were attenuated when SES indicators were included as covariates (Figure 6; Supplemental Tables 5–7). Epigenetic-*g* was positively and modestly associated with family-level SES in the sociodemographically diverse TTP (Epigenetic-*g* at T1 with parental SES: *r* = .08, 95% CI [.02, .14], *p* = .004, *n* = 1,130) and the larger GenR cord-blood Epigenetic-*g* (maternal education: *r* = .09, 95% CI [.05, .13], *p* < .001, *n* = 2,225). The association was not significantly different from zero in the smaller GenR venous-blood sample (*p* ≥ .130, *n* ≤ 471; see Figure 2) and in TwinLife, in which higher SES families are overrepresented (*p* ≥ .200, *n* ≤ 1,031; Frach et al., 2026).

Overall, adding SES covariates yielded small-to-moderate attenuation of biomarker-phenotype associations, with a stronger impact on Epigenetic-*g* than the PGIs. In TTP, Epigenetic-*g* coefficients declined by roughly 7–50% (from βs ≈ .18–.34 to βs ≈ .15– .22) while remaining largely robust after covariate adjustment across most cognitive and academic outcomes. In other cohorts, the comparatively smaller Epigenetic-*g* estimates were attenuated toward zero and no longer consistently survived FDR correction (with the exception that FFCWS estimates remained evident when adjusting for neighborhood, but not parental, SES).

SES adjustment produced predominantly modest attenuation of PGI associations. For PGI-Education, coefficients were typically reduced by approximately 6%–29% in TTP and TwinLife (from βs ≈ .17–.41 to βs ≈ .05–.38), with somewhat larger proportional attenuation for GenR age-12 academic outcomes (approximately 36%–38%). For PGI-Cognition, attenuation was generally smaller and often near-negligible (usually 0%–23%; from βs ≈ .11–.32 to βs ≈ .11–.29).

## Discussion

Leveraging birth cohort, longitudinal, and twin study designs, we examined the developmental correlates of an epigenetic index for adult cognitive function (’Epigenetic-*g*’) in comparison to polygenic indices (PGIs) of adult cognition and educational attainment. We addressed four key gaps concerning the stability of Epigenetic[g, its overlap with PGIs, how Epigenetic[g and PGIs relate to cognitive growth, and the genetic and environmental sources of variation in these measures.

*First*, Epigenetic-*g* was more dynamic early in life and became moderately stable by adolescence. In the Generation R Study, Epigenetic-*g* increased on average from ages 6 to 10 (∼0.1 SD per year), with significant between-person variability in intra-individual change. At the same time, despite tissue discordance, cord blood Epigenetic-*g* at birth was moderately associated with Epigenetic-*g* measured in venous blood at ages 6 and 10 (*r* ∼ 0.4). Across adolescence and into early adulthood in the Texas Twins and TwinLife studies, both the average level of Epigenetic-*g* and its rank-order stability increased, while intra-individual differences in change were minimal. Together, these patterns suggest that a substantial proportion of variability in Epigenetic-*g* emerges before early adolescence and is then carried forward with increasing stability across childhood and adolescence. This is consistent with epigenomic evidence that DNA methylation shapes prenatal tissue and organ differentiation and early childhood development (Mulder et al., 2021; Smith & Meissner, 2013).

*Second*, Epigenetic-*g* and PGIs were uncorrelated and accounted for distinct variance in children’s cognitive and academic performance. This is true even though our twin analyses indicated that a substantial amount of the variance in Epigenetic-*g* and its association with cognition is attributable to genetic variation (heritability of ∼60%). Our results mirror findings in adults reporting a comparable lack of association between polygenic and epigenetic indices of body mass index, alcohol consumption, and smoking (Hamilton et al., 2019; McCartney et al., 2018). It is consistent with the notion that epigenetic indices capture direct genetic regulation of methylation (i.e., meQTLs), non-additive genetic effects (e.g., gene-gene interactions), and specific gene-environment interactions (e.g., gene-tissue) that contribute to twin heritability, but are not captured by current PGIs (Cheesman et al., 2023; Villicaña & Bell, 2021). Thus, trait-matched epigenetic and polygenic biomarkers capture largely distinct genomic variation.

Although PGIs showed more robust associations with cognition and academic achievement across cohorts and measures, Epigenetic-*g’s* effect sizes were of a similar magnitude and accounted for distinct variation. Across the four cohorts and tissue types, children with higher Epigenetic-*g* had higher non-verbal cognitive test performance. In the Texas Twin Project, which has a particularly extensive battery of cognitive and academic measures compared to other cohorts, the combined variance accounted for by Epigenetic-*g* and PGIs was ∼22% in state-mandated reading tests, of which ∼9% was unique to Epigenetic-*g*. This result is particularly striking because Epigenetic-*g* was developed using a discovery sample several orders of magnitude smaller than those for PGIs, and Epigenetic-*g* only has a moderate cross-tissue correlation across blood and saliva measured in the same individuals (ICC = 0.69; Zarandooz & Raffington, 2025). Therefore, at birth, a child may have a low PGI of cognition, but a high cord-blood derived Epigenetic-*g*, and both are associated with their subsequent performance on cognitive tests relative to other children.

*Third*, unlike PGIs, Epigenetic-*g* was not associated with longitudinal cognitive growth across late childhood and adolescence. Children with higher PGIs were also more likely to read outside school and to be placed in more advanced academic tracks. Together with prior findings that literacy engagement and academic motivation partly account for the strengthening PGI–education links over development (Malanchini et al., 2017, 2024), this pattern is consistent with the view that PGI associations partly reflect ongoing person–environment transactions through which inherited differences are amplified over time. By contrast, Epigenetic-*g* appears more informative about stable or accumulated differences in performance than about later developmental change. These conclusions should be interpreted cautiously, as reduced statistical power in the smaller epigenetic subsample may have obscured such associations. Future research could explore whether novel epigenetic indices of adult or child cognition generated in larger discovery samples possess the measurement precision to detect individual differences in cognitive development.

*Fourth*, beyond additive genetic influences, variance in Epigenetic-*g* was predominantly attributable to processes unique to individual children rather than to environmental factors shared within families. Similar to PGIs, Epigenetic-*g* effect sizes were attenuated in within-family comparisons, with attenuation more pronounced for academic outcomes than for cognition. Further supporting this interpretation, associations between Epigenetic-*g* and family socioeconomic status were modest, and adjusting for socioeconomic status left cognitive associations largely unchanged while more clearly attenuating academic associations. Together, these patterns suggest that genomic associations with academic performance are more strongly shaped by family-shared environmental factors (for example, parental education and schooling) than are associations with cognitive performance — and that much of the epigenetic variation is unique to each child.

Our analyses of monozygotic twins further revealed that this unique variation in Epigenetic-*g* is meaningful: Differences between monozygotic twins in their Epigenetic-*g* were associated with differences in their fluid cognitive performance in both laboratory (i.e., perceptual reasoning and calculation skills) and school settings (i.e., math). Such differences may arise from stochastic developmental processes or from early environmental exposures that diverge between co-twins, including differences in intrauterine position, birth weight, or child-specific stress during sensitive periods of neurodevelopment. This substantial nonshared environmental variation, estimated at approximately 40%, may reflect ‘real but random’ biological and psychological effects on the epigenome (Plomin, 2024). Studies in adolescents and adults have suggested that such idiosyncratic processes in the epigenome, also referred to as ‘epigenetic drift’, may be critically involved in biological aging and health (Horvath, 2013; Horvath & Raj, 2018; Instinske, Schowe, et al., 2026; Kuznetsov et al., 2025). Epigenetic indices therefore offer a tractable way to study these processes in relation to behavior and health.

This study has several limitations. Some analyses were constrained by modest sample sizes, particularly in sibling contrasts and longitudinal epigenetic analyses, requiring cautious interpretation especially of null results. Additionally, the discovery samples used to generate the genomic indices lacked sociodemographic diversity and originated in blood samples collected from adults (McCartney et al., 2022). This limits the portability of the indices to pediatric or sociodemographically diverse populations, as epigenetic patterns are known to vary by age, ancestry, and social context, potentially introducing bias when applied to samples that differ systematically from those used in discovery. Finally, several cognitive and academic phenotypes were measured with brief, heterogeneous instruments across cohorts and ages, which limits construct coverage and cross-cohort comparability. Future work may benefit from leveraging larger and more diverse discovery samples, harmonized longitudinal cognitive measures, and repeated tissue-matched DNAm collections to improve portability, precision, and interpretability.

In conclusion, our findings highlight development as the dynamic interface between genotype and environment that associates variation in DNA and DNA methylation with adult cognition and educational attainment. We find that Epigenetic-*g* captures complementary genomic signal that is largely independent of PGIs and detectable from early life. Thus, epigenetic indices derived from peripheral tissues not only yield widely used estimates of biological aging, known as ‘epigenetic clocks’, but also capture biological signals associated with cognitive processes that are partially independent of stable DNA sequence – as evidenced by associations within monozygotic twin pairs. These observations align with extended models of heritability that view development, and the epigenome, not merely as an interpreter of genes but as an active contributor to phenotypic variation (Freund et al., 2013; Lala, 2024; Molenaar et al., 1993). Given that genomic biomarkers are increasingly applied in clinical and policy contexts, including embryo screening (Capalbo et al., 2024; Turley et al., 2021), understanding their developmental foundations is especially pressing. As epigenetic discovery samples expand in size, ancestral diversity, and developmental coverage, epigenetic indices may increasingly complement PGIs in characterizing the genomic architecture of cognitive and academic development within population-level research.

## Methods

### Participants

#### Texas Twin Project (TTP)

The TTP is an ongoing population-representative longitudinal study investigating children and adolescents in the greater metropolitan areas of Austin, Texas and Houston, USA (Harden et al., 2013). Polygenic analyses were restricted to individuals whose DNA-based ancestries closely align with the GWAS discovery sample’s ancestries (i.e., European ancestries) to avoid the risk of confounding due to population stratification (*n* = 896). DNAm data were available for *n* = 1,823 children and adolescents (*M_age_* = 13.72, *SD_age_* = 2.88, 50.7% female, 34.6% monozygotic twins, 58.9% dizygotic twins, 59.5% identified as White, 10.7% as Hispanic/Latinx-only, 10.4% as Black/African-American, 8.5% as Asian, and 7.8% as Latinx-White).

#### Future of Families and Child Wellbeing Study (FFCWS)

The FFCWS follows a sample of 4,898 children born in large US cities during 1998 – 2000. FFCWS oversampled children born to unmarried parents and interviewed parents at birth and ages 1, 3, 5, 9 and 15. During home visits, saliva DNA was collected with the Illumina 450K and EPIC methylation arrays with ages 9 (*n* = 1971) and 15 years (*n* = 1974) assayed on the same plate (for further information on DNA preprocessing see Supplemental Material). DNAm study participants self-identified race/ethnicity defined by study protocol as African-American/Black only (*n* = 901, 47%), ‘Other’ (*n* = 52, 3%), Hispanic/Latinx (*n* = 511, 26%), Multiracial (*n* = 99, 5%), White (*n* = 366, 19%). The University of Michigan and Princeton University Institutional Review board granted ethical approval. Informed written consent was obtained from all participants and study participants’ legal guardians. All methods were performed in accordance with the relevant guidelines and regulations.

#### The Generation R Study (GenR)

GenR is a prospective population-based birth cohort. Pregnant women residing in the study area of Rotterdam, The Netherlands, with an expected delivery date between April 2002 and January 2006 were invited to enroll and 9,749 live-born children were included. A more extensive description of the study is detailed elsewhere (Kooijman et al., 2016). Genotyping was performed using Illumina 610K Quad Chip or Illumina 660k Quad Chip, based on cord-blood samples at birth or peripheral blood samples in childhood. Genotype data were imputed to the 1000 Genomes reference panel. For more information, (see Medina-Gomez et al., 2015). DNAm was measured on a genome-wide level in a subgroup of Dutch children, using the Illumina Infinium MethylationEPIC v1.0 (EPICv1) BeadChip and Illumina Infinium HumanMethylation450 (450K) BeadChip (Illumina Inc., San Diego, USA). EPICv1 was only applied at birth, in cord-blood, in 1,115 samples. 450K was applied in a separate set of cord-blood samples of 1339 children, blood samples in 469 children aged 6 years and blood samples in 425 children aged 10 years. Preparation and normalization of the data were performed according to the CPACOR workflow (Lehne et al., 2015), using the software package R. The data were quantile normalized, and DNAm betas were winsorized (median +/- 3 IQR) to the nearest point for each DNAm site to reduce the influence of potential outliers. The Generation R Study is conducted in accordance with the World Medical Association Declaration of Helsinki and has been approved by the Medical Ethics Committee of Erasmus MC, University Medical Center Rotterdam. Written informed consent was obtained for all participants.

#### The TwinLife study

The German Twin Family Panel (TwinLife) is a representative, cross-sequential longitudinal study of twins and family members (Diewald et al., 2025; Rohm et al., 2023). The genome-wide genotyping in TwinLife is ongoing. As of now, genotyping was performed for a subset of *n* = 5,881 TwinLife participants. PGI analyses were restricted to participants with the GWAS discovery sample’s ancestries (i.e., European ancestries) to avoid the risk of confounding due to population stratification. This results in a post-QC genome-wide genotype and PGI dataset for *n* = 5,432 twins, their siblings and parents (55% female).

Based on the availability of funds, DNA samples of *n* = 1,064 twins (M_age_= 15.86, SD_age_ = 6.05 at baseline and M_age_= 18.16, SD_age_ = 6.07 at follow-up, 53% female, 53% monozygotic twins, 47% dizygotic twins) were selected for DNAm analyses. The inclusion criteria were as follows: 1) valid saliva samples of twin children and adolescents that passed the genotyping and PGI calculation quality control steps, 2) participants that are twins and participated in the third face-to-face interview, as well as the second and third COVID-19 supplementary survey of the TwinLife study, 3) inclusion of all participants with two complete saliva samples from complete pairs of twins. The remainder of the available array slots were then filled with twins with two complete saliva samples but no eligible samples available for their corresponding twin. The TwinLife study was reviewed and ethically approved by the German Psychological Society (Deutsche Gesellschaft für Psychologie; protocol number: RR 11.2009). The Ethics Committee of the Medical Faculty of the University of Bonn (No. 113/18) reviewed and approved the protocols for the molecular genetic analyses based on saliva samples. Informed written consent (for molecular genetic data) and informed verbal consent (for phenotypic data) were obtained from all participants and the participant’s legal guardian (for minors).

**Table 1.**
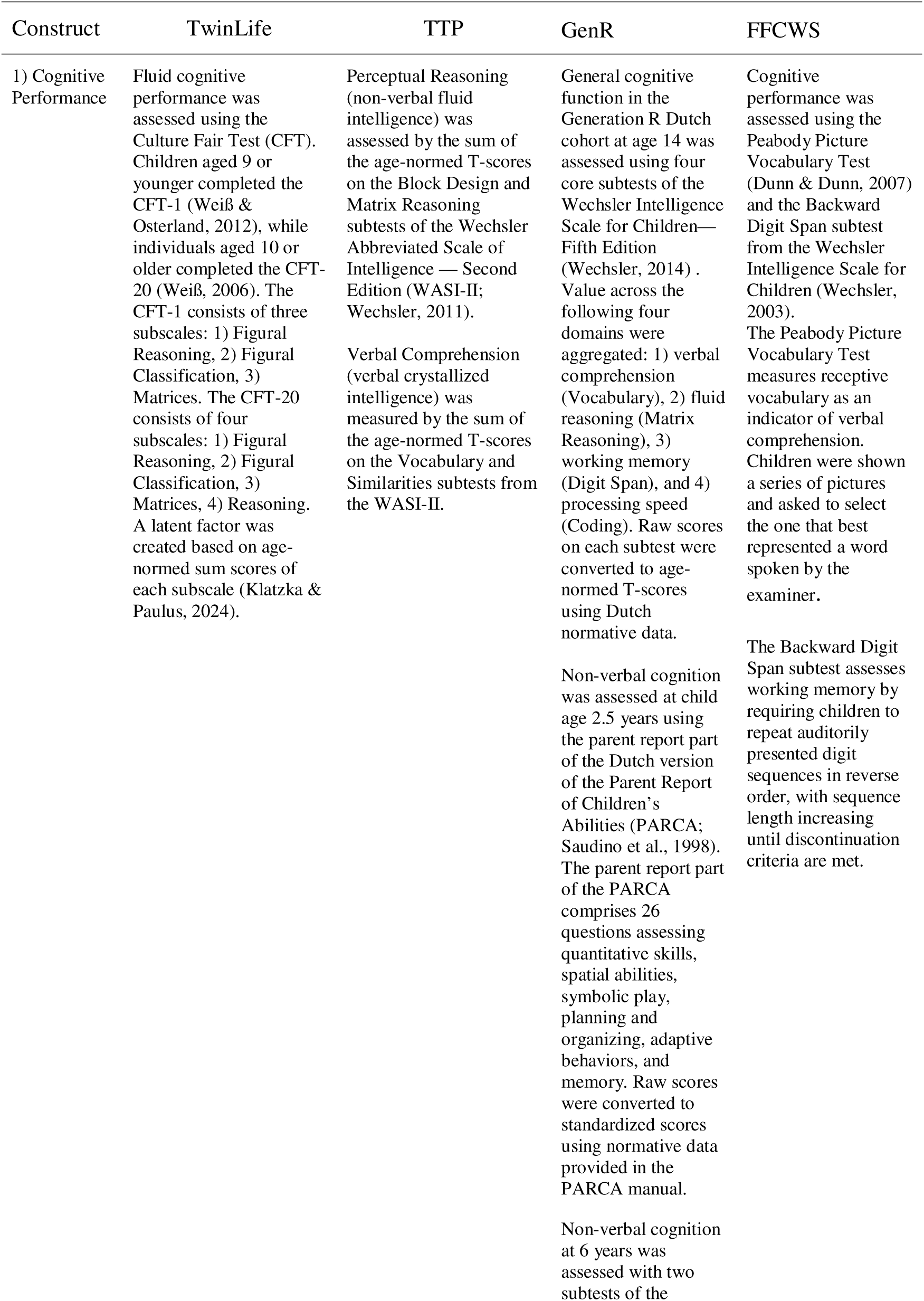
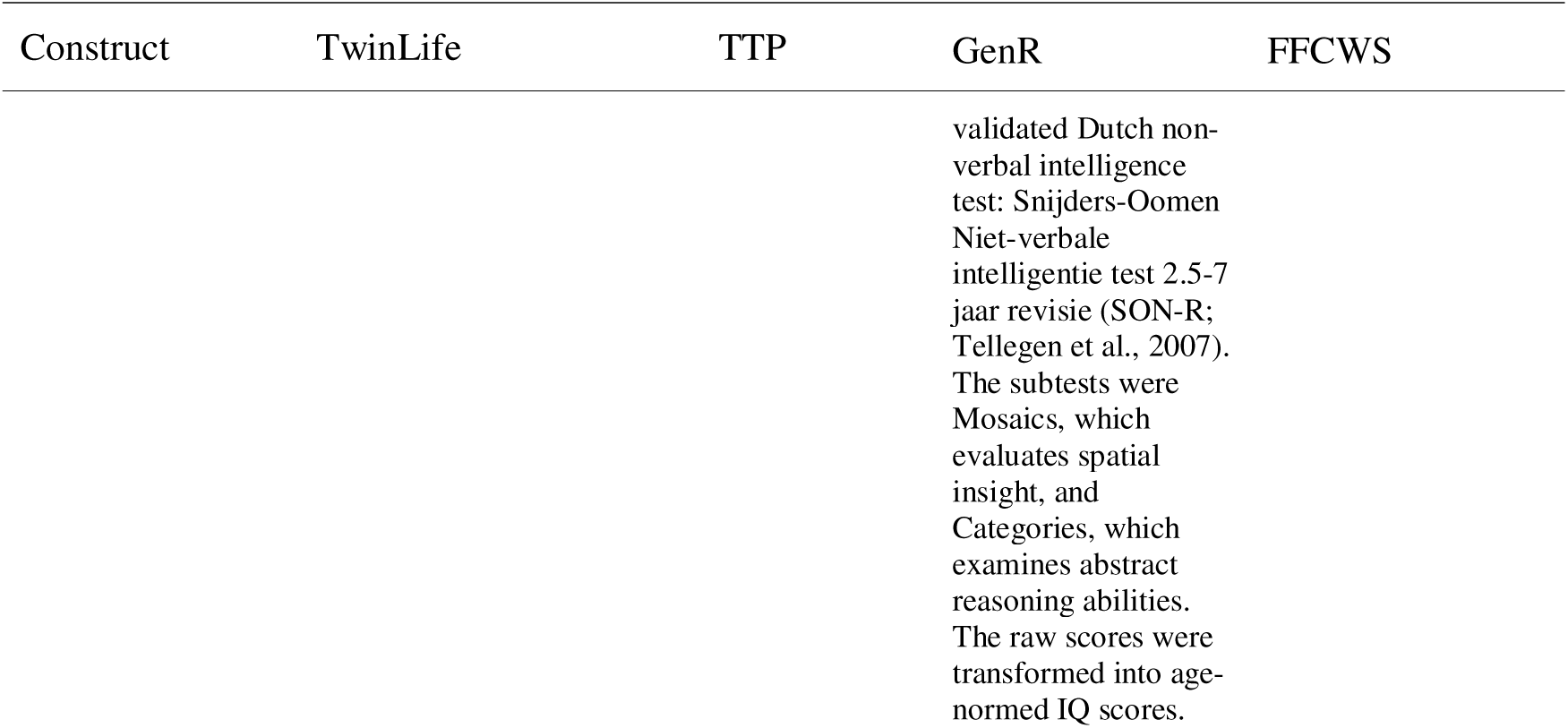
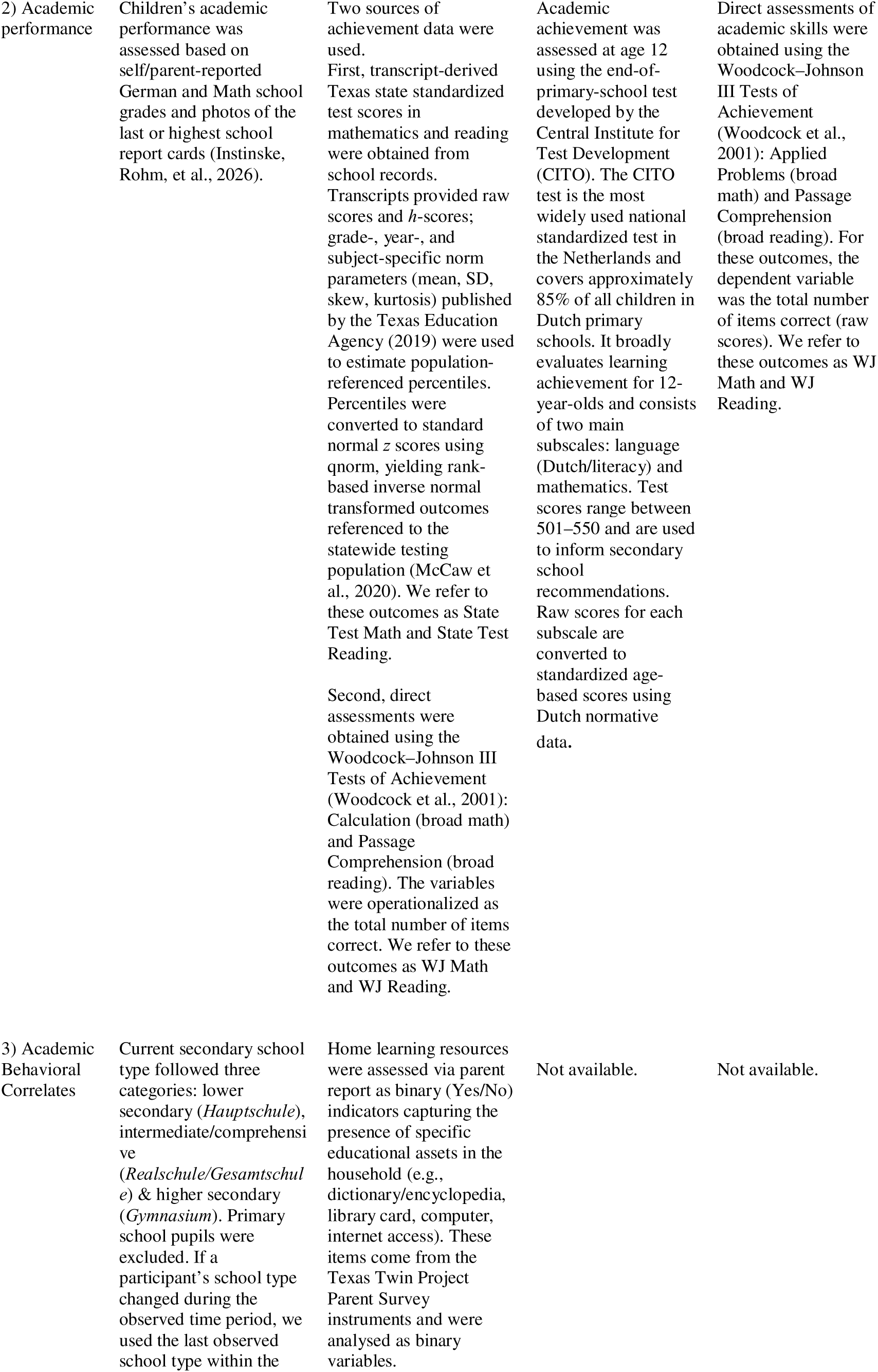
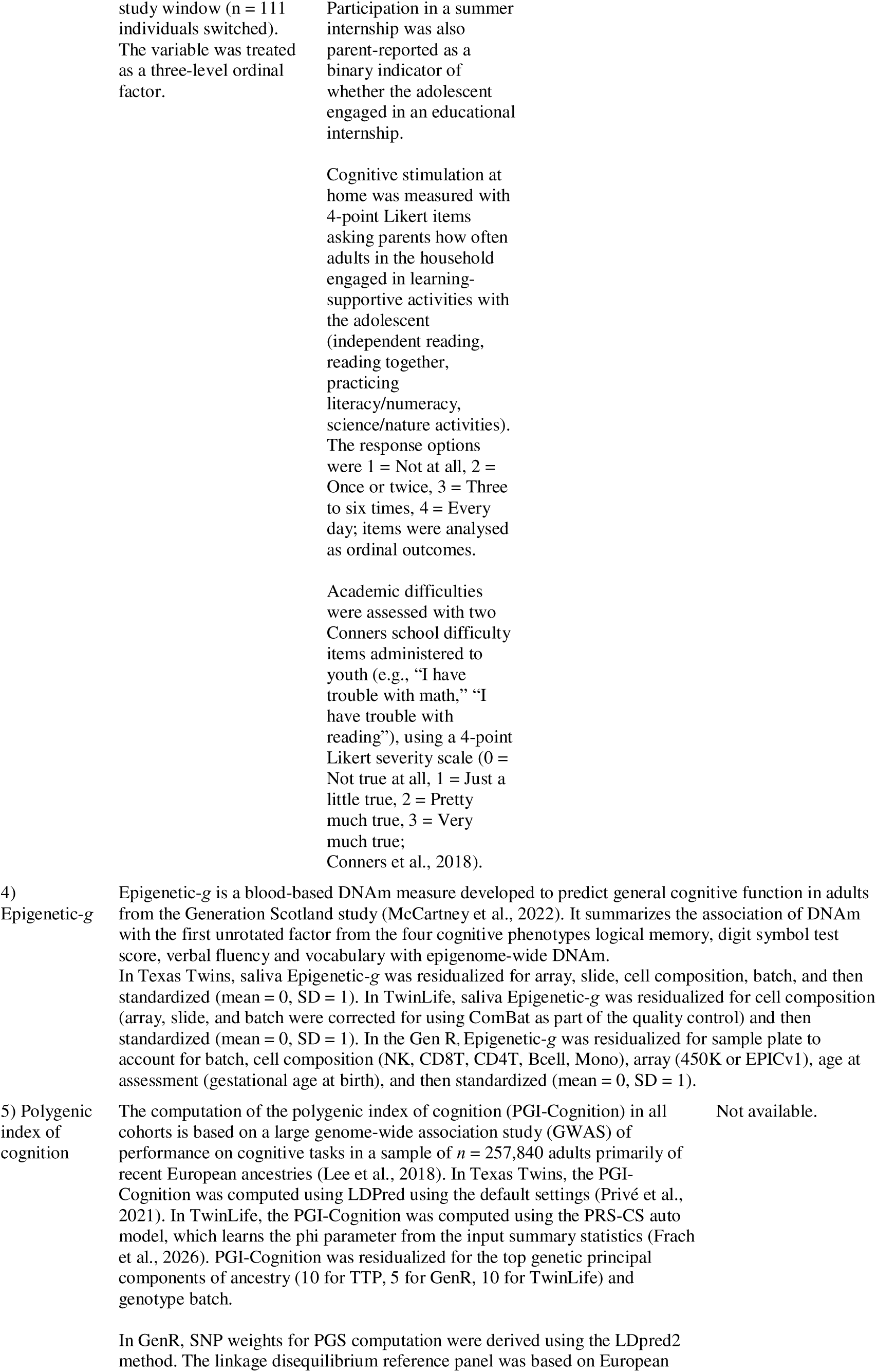
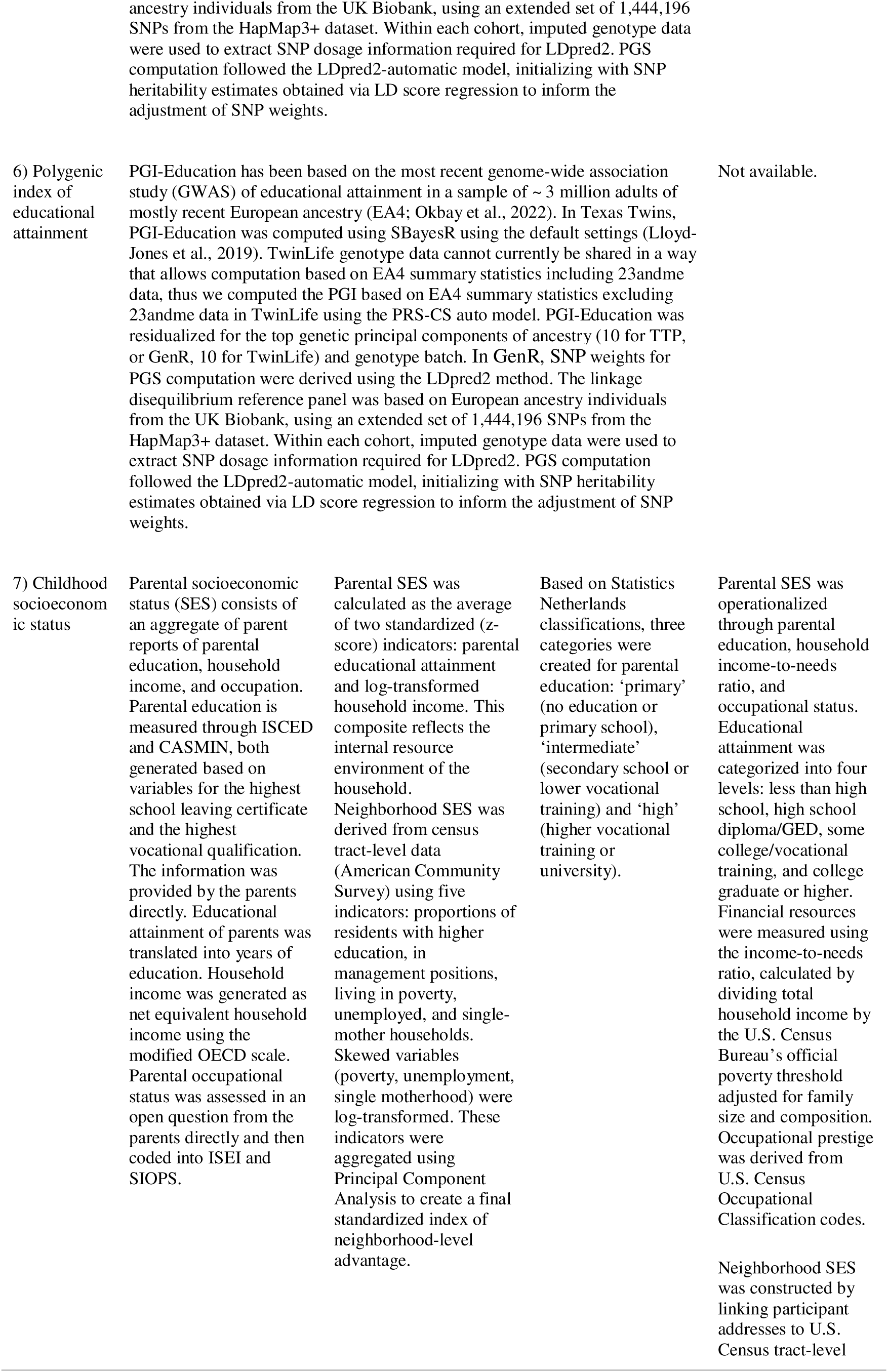
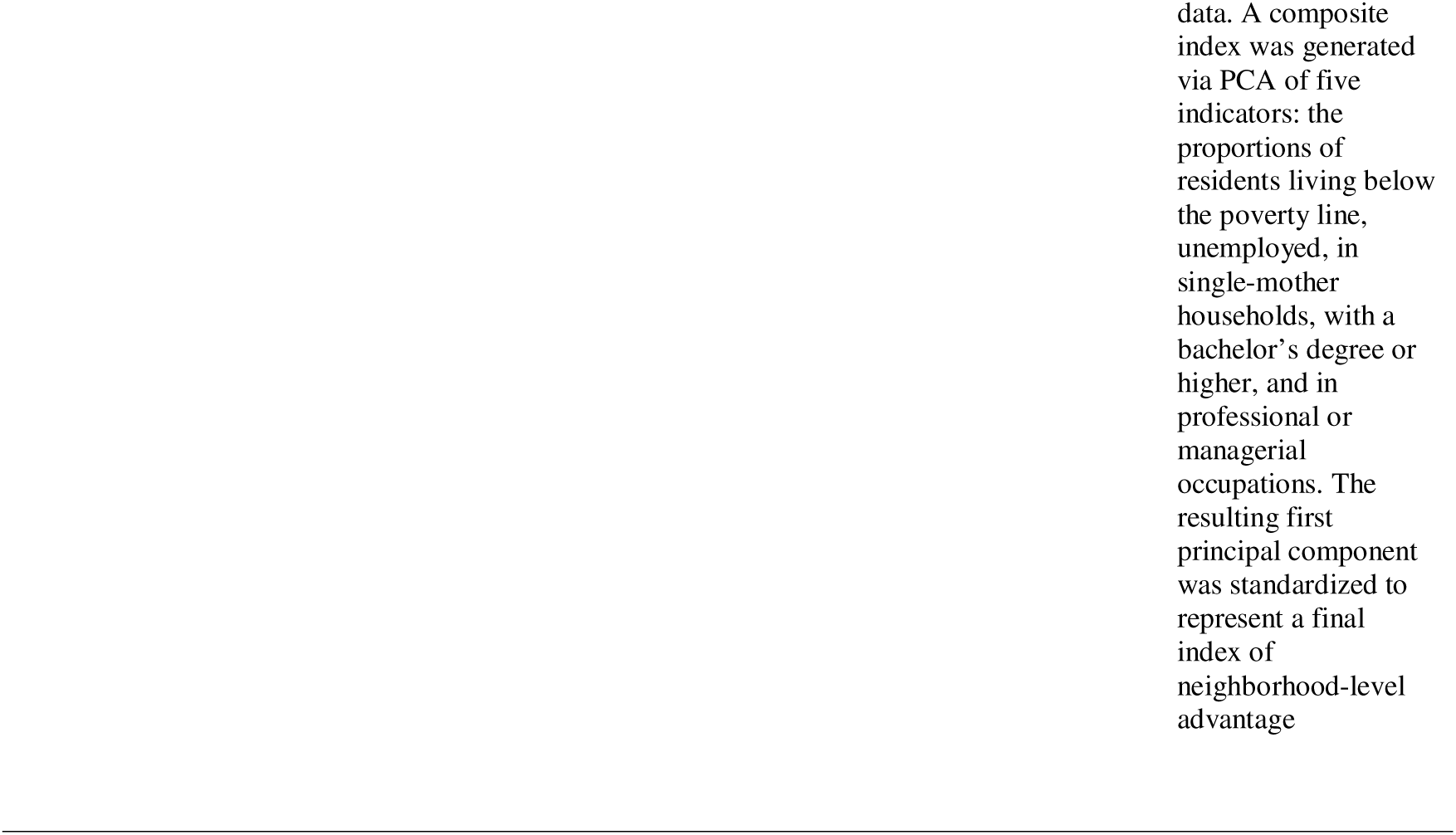
Description of main variables of interest across cohorts.

## Measures

### Statistical Analyses

All analyses were conducted in R and Mplus 8.8 (Muthén & Muthén, 2017; R Core Team, 2024). We used two-level random-effects models with repeated observations nested within individuals and cluster-robust standard errors for families. We conducted five sets of statistical analyses.

1. For time-variant variables from the TTP and TwinLife, we fit latent growth-curve models (LGCMs) using an accelerated longitudinal (cohort-sequential) specification that leverages overlapping age coverage across participants (Duncan et al., 2006; McArdle & Bell, 2000). These models estimate each individual’s baseline level (latent intercept) and rate of within-person change (latent slope), allowing us to model developmental trajectories across childhood and adolescence. Curvature was modeled by including a within-person quadratic age term in TTP and a logarithmic age term in TwinLife, as these provided best model fit within each cohort. Age (in years; decades in TTP for convergence) was centered to the first DNAm assessment (7.8 years). The two postnatal waves of DNAm in GenR were analysed using latent difference score models (LDSMs), which estimate the average within-person change between two measurement occasions as well as interindividual differences in that change.
2. Using two-level latent intercept models aligned with the growth specifications in (1), we regressed latent intercepts of cognitive performance and academic achievement on (a) PGI-Education, (b) PGI-Cognition, and (c) the Epigenetic-*g* intercept, first separately and then jointly to decompose unique versus shared variance. In FFCWS, cross-sectional associations were estimated by regressing each outcome on Epigenetic-*g* at the first available DNAm assessment (Year 9).
3. In cohorts with repeated cognitive or academic assessments (TTP, TwinLife), we fit two-level LGCMs in line with (1), where age (centered and scaled within cohort) defined within-person change, and time-invariant correlates (PGI-Education, PGI-Cognition, or the Epigenetic-*g* intercept) predicted the latent intercepts and slopes, adjusted for sex.
4. To probe family-level confounding, we estimated (i) pair-demeaned regressions (family fixed effects) in twins/siblings and (ii) univariate and bivariate Cholesky models partitioning additive genetic (A), shared environmental (C), and non-shared environmental (E) sources of variance in Epigenetic-*g* and its covariance with cognition/achievement. As a complementary analysis, absolute MZ within-pair differences in Epigenetic-*g* were related to absolute within-pair outcome differences.
5. Childhood SES was modeled as an exogenous covariate. We report attenuated associations when adding SES to models in 2–3 and describe SES–biomarker correlations.

Across analyses, variables were z-standardized. For FFCWS, only unstandardized regression coefficients were available. Since all predictors were z-standardized, we approximated standardized coefficients by dividing unstandardized estimates by the published raw-score standard deviations of each outcome (Future Families and Child Wellbeing Study, 2011). This procedure permits scale-comparable effect sizes across cohorts given the constraints of the available data. Missingness was handled via FIML under MAR; estimation used MLR with family clustering. Multiple testing was controlled using the Benjamini–Hochberg FDR procedure within each analysis set and outcome domain. Full model specifications are provided in the Supplementary Methods. The code will be publicly available upon publication at https://github.com/Biosocial/EpiG_PGI_Cognition.

## Data Availability Statement

This study utilizes data from the Future Families and Child Wellbeing Study (FFCWS). Researchers interested in accessing the data can find detailed information on the FFCWS website at https://ffcws.princeton.edu, where additional documentation is available at https://ffcws.princeton.edu/documentation. Publicly available, de-identified data from the previous six waves of data collection can be accessed through the Princeton University Office of Population Research (OPR) Data Archive at https://ffcws.princeton.edu/data-and-documentation/public-data-documentation. For researchers requiring restricted-use contract data, access is granted only to those who agree to the terms outlined in the FFCWS Contract Data License. Further details on obtaining restricted data can be found at https://ffcws.princeton.edu/restricted.

In the TTP access to polygenic and phenotype data is available only under a restricted data use agreement with the University of Texas at Austin. Applicants must submit an IRB-approved research proposal and a data security plan; instructions and the application portal are provided by the TTP/UT Austin team. Details on procedures and the application link are available via the Population Research Center’s restricted-use application page for TTP.

TwinLife survey data are curated and distributed via the GESIS Data Catalogue and are available for academic research and teaching upon completion of the GESIS/TwinLife Data Use Agreement. Comprehensive study documentation, codebooks, and a short guide are publicly available (https://www.twin-life.de/documentation); instructions for requesting survey data are provided in the documentation and catalogue entry. At present, the genetic data cannot be made available via the GESIS because of ethical reasons regarding sensitive data and can only be accessed within the framework of a scientific collaboration with the TwinLife team. In the near future, the genetic data will be made available through the Research Data Centre at Bielefeld University.

In the Generation R Study, individual-level data are not publicly available due to participant privacy and governance restrictions. The study operates an open-collaboration policy: external researchers may request collaboration by submitting a proposal to the Generation R Management Team (contacted via the study leadership at Erasmus MC). Proposals are reviewed for scientific merit, overlap with ongoing projects, logistics, and required resources; approved projects proceed under institutional ethical approvals.

## Author contributions

Conceptualization: DF, LR, JI, CK Methodology: DF, LR

Data Collection: BM, EMTD, KPH, MMN, CKLD, AJF

Data Curation: AMS, PTT, LP, DF, CD, JI, CK

Analysis: DF, LP, IS, CM, JW

Writing - Original Draft: DF, LR

Writing - Review & Editing: All authors

Supervision: LR, EMTD

Funding Acquisition: LR, CM, EMTD, KPH, CCAM, EBB, AJF, MMN, FMS, CK, BM

## Acknowledgments

The content is solely the responsibility of the authors and does not necessarily represent the official views of the funders. We acknowledge access to the bonna cluster at the University of Bonn and the support provided by the HPC@HRZ team. This research was conducted while CCAM was a Hevolution/AFAR New Investigator Awardee in Aging Biology and Geroscience Research. The Generation R Study is conducted by Erasmus MC, University Medical Center Rotterdam, in close collaboration with the School of Law and Faculty of Social Sciences of Erasmus University Rotterdam, the Municipal Health Service Rotterdam area, Rotterdam, the Rotterdam Homecare Foundation, Rotterdam, and the Stichting Trombosedienst & Artsenlaboratorium Rijnmond (STAR-MDC), Rotterdam. We gratefully acknowledge the contribution of children and parents, general practitioners, hospitals, midwives, and pharmacies in Rotterdam. The study protocol was approved by the Medical Ethical Committee of Erasmus MC, Rotterdam. The generation and management of the GWAS genotype data and the DNA methylation array data for the Generation R Study were carried out at the Human Genomics Facility (HuGe-F), housed within the Laboratory for Population Genomics of the Department of Internal Medicine at Erasmus MC. We thank M. Verbiest, M. Jhamai, S. Higgins, M. Verkerk, and L. Stolk for their help in creating the EWAS database. We thank A. Teumer for work on EWAS quality control and normalization. We thank P. Arp, K. Estrada, M. Jhamai, M. Ganesh, G. van Dijk, M. Verkerk, L. Herrera, M. Peters, and L. Broer for their help in creating, managing, and quality-controlling the GWAS database. MM is supported by a Jacobs Foundation Research Fellowship (#2024-1533-00) and UK Medical Research Council Grant UKRI1506. KPH was supported by grants R01HD092548 and R01HD114724 from the Eunice Kennedy Shriver National Institute of Child Health and Human Development (NICHD). KPH and EMTD are Faculty Research Associates of the Population Research Center at the University of Texas at Austin, which is supported by grant P2CHD042849 from NICHD.

## Funding

Max Planck Society (LR, EBB, UL)

National Institutes of Health grant R01HD114724 (LR, KPH)

National Institutes of Health grant R01HD092548 (KPH, EMTD)

National Institutes of Health grant R01AG073593 (EMTD)

National Institutes of Health grant P2CHD042849 (KPH, EMTD)

Jacobs Foundation (LR, MM)

European Union grant #101073237 (ESSGN; LR)

European Union grant #101057529 (FAMILY; CC)

European Union grant #101057390 (HappyMums; IS, CC)

European Research Council grant #101039672 (TEMPO; IS, CC)

IMPRS Life (DF, LR, UL)

UK Medical Research Council Grant UKRI1506 (MM)

This project has received funding from the European Union’s Horizon Europe Research and Innovation Programme (STAGE, grant agreement No 101137146 [JF, CC], FAMILY, Grant Agreement No. 101057529 [CC]; HappyMums, Grant Agreement No. 101057390 [CC, IS]) and the European Research Council (TEMPO; Grant Agreement No. 101039672, [CC, IS]). Views and opinions expressed are however those of the author(s) only and do not necessarily reflect those of the European Union. Neither the European Union nor the granting authority can be held responsible for them.

The TwinLife study was funded by the German Research Foundation (DFG) grants 220286500 (CK, FMS, BM), 428902522 (MMN, FMS), and 458609264 (EB, AJF, CK, MMN, FMS).

## Notes

### Competing Interest Statement

The authors have declared no competing interest.

## References

Abdellaoui, A., Martin, H. C., Kolk, M., Rutherford, A., Muthukrishna, M., Tropf, F. C., Mills, M. C., Zietsch, B. P., Verweij, K. J. H., & Visscher, P. M. (2025). Socio-economic status is a social construct with heritable components and genetic consequences. Nature Human Behaviour, 9(5), 864–876. 10.1038/s41562-025-02150-4

Allegrini, A. G., Selzam, S., Rimfeld, K., von Stumm, S., Pingault, J. B., & Plomin, R. (2019). Genomic prediction of cognitive traits in childhood and adolescence. Molecular Psychiatry, 24(6), 819–827. 10.1038/s41380-019-0394-4

Beam, C. R., Turkheimer, E., Finkel, D., Levine, M. E., Zandi, E., Guterbock, T. M., Giangrande, E. J., Ryan, L., Pasquenza, N., & Davis, D. W. (2020). Midlife Study of the Louisville Twins: Connecting Cognitive Development to Biological and Cognitive Aging. Behavior Genetics, 50(2), 73–83. 10.1007/s10519-019-09983-6

Bertucci-Richter, E. M., Shealy, E. P., & Parrott, B. B. (2024). Epigenetic drift underlies epigenetic clock signals, but displays distinct responses to lifespan interventions, development, and cellular dedifferentiation. Aging, 16(2), 1002–1020. 10.18632/aging.205503

Capalbo, A., de Wert, G., Mertes, H., Klausner, L., Coonen, E., Spinella, F., Van de Velde, H., Viville, S., Sermon, K., Vermeulen, N., Lencz, T., & Carmi, S. (2024). Screening embryos for polygenic disease risk: A review of epidemiological, clinical, and ethical considerations. Human Reproduction Update, 30(5), 529–557. 10.1093/humupd/dmae012

Cheesman, R., Ayorech, Z., Eilertsen, E. M., & Ystrom, E. (2023). Why we need families in genomic research on developmental psychopathology. JCPP Advances, 3(1), e12138. 10.1002/jcv2.12138

Conners, C. K., Rzepa, S. R., Pitkanen, J., & Mears, S. (2018). Conners 3rd Edition (Conners 3; Conners 2008). In Encyclopedia of Clinical Neuropsychology (pp. 1–5). Springer, Cham. 10.1007/978-3-319-56782-2_1534-2

de Zeeuw, E. L., van Beijsterveldt, C. E. M., Glasner, T. J., Bartels, M., Ehli, E. A., Davies, G. E., Hudziak, J. J., Consortium, S. S. G. A., Rietveld, C. A., Groen-Blokhuis, M. M., Hottenga, J. J., de Geus, E. J. C., & Boomsma, D. I. (2014). Polygenic scores associated with educational attainment in adults predict educational achievement and ADHD symptoms in children. American Journal of Medical Genetics Part B: Neuropsychiatric Genetics, 165(6), 510–520. 10.1002/ajmg.b.32254

Deary, I. J., Weiss, A., & Batty, G. D. (2010). Intelligence and Personality as Predictors of Illness and Death: How Researchers in Differential Psychology and Chronic Disease Epidemiology Are Collaborating to Understand and Address Health Inequalities. Psychological Science in the Public Interest: A Journal of the American Psychological Society, 11(2), 53–79. 10.1177/1529100610387081

Demange, P. A., Malanchini, M., Mallard, T. T., Biroli, P., Cox, S. R., Grotzinger, A. D., Tucker-Drob, E. M., Abdellaoui, A., Arseneault, L., van Bergen, E., Boomsma, D. I., Caspi, A., Corcoran, D. L., Domingue, B. W., Harris, K. M., Ip, H. F., Mitchell, C., Moffitt, T. E., Poulton, R.,…Nivard, M. G. (2021). Investigating the genetic architecture of noncognitive skills using GWAS-by-subtraction. Nature Genetics, 53(1), Article 1. 10.1038/s41588-020-00754-2

deSteiguer, A. J., Raffington, L., Sabhlok, A., Tanksley, P., Tucker-Drob, E. M., & Harden, K. P. (2025). Stability of Aging- and Cognition-Related Methylation Profile Scores Across Two Waves in Children and Adolescents. Child Development, 96(3), 1189–1206. 10.1111/cdev.14239

Diewald, M., Kandler, C., Riemann, R., Spinath, F. M., Mönkediek, B., Andreas, A., Baier, T., Bartling, A., Baum, M. A., Deppe, M., Eichhorn, H., Eifler, E. F., Gottschling, J., Hahn, E., Hildebrandt, J., Hufer, A., Instinske, J., Kaempfert, M., Klatzka, C. H.,…Weigel, L. (2025). TwinLife. 10.4232/1.14531

Duncan, T. E., Duncan, S. C., & Strycker, L. A. (2006). An Introduction to Latent Variable Growth Curve Modeling: Concepts, Issues, and Applications (2nd ed.). Lawrence Erlbaum Associates Publishers.

Dunn, L. M., & Dunn, D. M. (2007). Peabody Picture Vocabulary Test (4th ed.).

Faul, J. D., Collins, S., Smith, T., Klopack, E. T., Mitchell, C., Crimmins, E. M., & Farina, M. P. (2025). Epigenetic g predicts cognitive aging and incident dementia in a diverse, nationally representative sample of older adults (p. 2025.12.23.25342931). medRxiv. 10.64898/2025.12.23.25342931

Frach, L., Disselkamp, C. K. L., Schowe, A. M., Andreas, A., Deppe, M., Instinske, J., Maj, C., Rohm, T., Ruks, M., Wiesmann, L. M., Kandler, C., Mönkediek, B., Spinath, F. M., Nöthen, M. M., Binder, E. B., Czamara, D., & Forstner, A. J. (2026). (Epi-)Genomic Data in the German TwinLife Study: TwinSNPs and TECS Cohort Profiles (p. 2026.02.20.704007). bioRxiv. 10.64898/2026.02.20.704007

Freund, J., Brandmaier, A. M., Lewejohann, L., Kirste, I., Kritzler, M., Krüger, A., Sachser, N., Lindenberger, U., & Kempermann, G. (2013). Emergence of individuality in genetically identical mice. Science, 340(6133), 756–759. 10.1126/science.1235294

Future Families and Child Wellbeing Study. (2011). Data User’s Guide for the Nine-Year Follow-Up Wave of the Future Families and Child Wellbeing Study. https://ffcws.princeton.edu/sites/g/files/toruqf4356/files/ff_public_guide_9.pdf

Hamilton, O. K. L., Zhang, Q., McRae, A. F., Walker, R. M., Morris, S. W., Redmond, P., Campbell, A., Murray, A. D., Porteous, D. J., Evans, K. L., McIntosh, A. M., Deary, I. J., & Marioni, R. E. (2019). An epigenetic score for BMI based on DNA methylation correlates with poor physical health and major disease in the Lothian Birth Cohort. International Journal of Obesity (2005), 43(9), 1795–1802. 10.1038/s41366-018-0262-3

Harden, K. P., Domingue, B. W., Belsky, D. W., Boardman, J. D., Crosnoe, R., Malanchini, M., Nivard, M., Tucker-Drob, E. M., & Harris, K. M. (2020). Genetic associations with mathematics tracking and persistence in secondary school. Npj Science of Learning, 5(1), 1. 10.1038/s41539-020-0060-2

Harden, K. P., Tucker-Drob, E. M., & Tackett, J. L. (2013). The Texas Twin Project. Twin Research and Human Genetics, 16(1), 385–390. 10.1017/thg.2012.97

Horvath, S. (2013). DNA methylation age of human tissues and cell types. Genome Biology, 14(10), 3156. 10.1186/gb-2013-14-10-r115

Horvath, S., & Raj, K. (2018). DNA methylation-based biomarkers and the epigenetic clock theory of ageing. Nature Reviews Genetics, 19(6), 371–384. 10.1038/s41576-018-0004-3

Instinske, J., Rohm, T., Mattheus, S., Starr, A., & Riemann, R. (2026). Documentation TwinLife Data: Report Cards (TwinLife Technical Report Series 04) [Report]. Universität Bielefeld / Universität Bremen / Universität des Saarlandes. https://pub.uni-bielefeld.de/record/3013285

Instinske, J., Schowe, A. M., Czamara, D., Kuznetsov, D. V., Mönkediek, B., & Kandler, C. (2026). Epigenetic aging and personality differences: Latent change analyses of twin data. European Journal of Personality, 08902070261418426. 10.1177/08902070261418426

Klatzka, C. H., & Paulus, L. (2024). Documentation TwinLife Data: Cognitive Abilities F2F1, F2F2, & F2F4 v.2.0.0 (TwinLife Technical Report Series 02) [Report]. Universität Bielefeld / Universität Bremen / Universität des Saarlandes. https://pub.uni-bielefeld.de/download/2990575/2990576/TwinLife_TR_02_v2-0-0.pdf

Kooijman, M. N., Kruithof, C. J., van Duijn, C. M., Duijts, L., Franco, O. H., van IJzendoorn, M. H., de Jongste, J. C., Klaver, C. C. W., van der Lugt, A., Mackenbach, J. P., Moll, H. A., Peeters, R. P., Raat, H., Rings, E. H. H. M., Rivadeneira, F., van der Schroeff, M. P., Steegers, E. A. P., Tiemeier, H., Uitterlinden, A. G.,…Jaddoe, V. W. V. (2016). The Generation R Study: Design and cohort update 2017. European Journal of Epidemiology, 31(12), 1243–1264. 10.1007/s10654-016-0224-9

Kuznetsov, D. V., Liu, Y., Schowe, A. M., Czamara, D., Instinske, J., Disselkamp, C. K. L., Nöthen, M. M., Spinath, F. M., Binder, E. B., Diewald, M., Forstner, A. J., Kandler, C., & Mönkediek, B. (2025). Genetic and environmental contributions to epigenetic aging across adolescence and young adulthood. Clinical Epigenetics, 17(1), 78. 10.1186/s13148-025-01880-6

Lala, K. (2024). Evolution evolving: The developmental origins of adaptation and biodiversity. Princeton University Press.

Lee, J. J., Wedow, R., Okbay, A., Kong, E., Maghzian, O., Zacher, M., Nguyen-Viet, T. A., Bowers, P., Sidorenko, J., Karlsson Linnér, R., Fontana, M. A., Kundu, T., Lee, C., Li, H., Li, R., Royer, R., Timshel, P. N., Walters, R. K., Willoughby, E. A.,…Cesarini, D. (2018). Gene discovery and polygenic prediction from a genome-wide association study of educational attainment in 1.1 million individuals. Nature Genetics, 50(8), Article 8. 10.1038/s41588-018-0147-3

Lehne, B., Drong, A. W., Loh, M., Zhang, W., Scott, W. R., Tan, S.-T., Afzal, U., Scott, J., Jarvelin, M.-R., Elliott, P., McCarthy, M. I., Kooner, J. S., & Chambers, J. C. (2015). A coherent approach for analysis of the Illumina HumanMethylation450 BeadChip improves data quality and performance in epigenome-wide association studies. Genome Biology, 16(1), 37. 10.1186/s13059-015-0600-x

Lloyd-Jones, L. R., Zeng, J., Sidorenko, J., Yengo, L., Moser, G., Kemper, K. E., Wang, H., Zheng, Z., Magi, R., Esko, T., Metspalu, A., Wray, N. R., Goddard, M. E., Yang, J., & Visscher, P. M. (2019). Improved polygenic prediction by Bayesian multiple regression on summary statistics. Nature Communications, 10(1), 5086. 10.1038/s41467-019-12653-0

Malanchini, M., Allegrini, A. G., Nivard, M. G., Biroli, P., Rimfeld, K., Cheesman, R., von Stumm, S., Demange, P. A., van Bergen, E., Grotzinger, A. D., Raffington, L., De la Fuente, J., Pingault, J.-B., Tucker-Drob, E. M., Harden, K. P., & Plomin, R. (2024). Genetic associations between non-cognitive skills and academic achievement over development. Nature Human Behaviour, 1–13. 10.1038/s41562-024-01967-9

Malanchini, M., Wang, Z., Voronin, I., Schenker, V. J., Plomin, R., Petrill, S. A., & Kovas, Y. (2017). Reading self-perceived ability, enjoyment and achievement: A genetically informative study of their reciprocal links over time. Developmental Psychology, 53(4), 698–712. 10.1037/dev0000209

McArdle, J. J., & Bell, R. Q. (2000). An introduction to latent growth models for developmental data analysis. Methods in Psychological Research, 5(2), 69–107.

McCartney, D. L., Hillary, R. F., Conole, E. L. S., Banos, D. T., Gadd, D. A., Walker, R. M., Nangle, C., Flaig, R., Campbell, A., Murray, A. D., Maniega, S. M., Valdés-Hernández, M. del C., Harris, M. A., Bastin, M. E., Wardlaw, J. M., Harris, S. E., Porteous, D. J., Tucker-Drob, E. M., McIntosh, A. M.,…Marioni, R. E. (2022). Blood-based epigenome-wide analyses of cognitive abilities. Genome Biology, 23, 26. 10.1186/s13059-021-02596-5

McCartney, D. L., Hillary, R. F., Stevenson, A. J., Ritchie, S. J., Walker, R. M., Zhang, Q., Morris, S. W., Bermingham, M. L., Campbell, A., Murray, A. D., Whalley, H. C., Gale, C. R., Porteous, D. J., Haley, C. S., McRae, A. F., Wray, N. R., Visscher, P. M., McIntosh, A. M., Evans, K. L.,…Marioni, R. E. (2018). Epigenetic prediction of complex traits and death. Genome Biology, 19(1), 136. 10.1186/s13059-018-1514-1

McCaw, Z. R., Lane, J. M., Saxena, R., Redline, S., & Lin, X. (2020). Operating Characteristics of the Rank-Based Inverse Normal Transformation for Quantitative Trait Analysis in Genome-Wide Association Studies. Biometrics, 76(4), 1262–1272. 10.1111/biom.13214

Medina-Gomez, C., Felix, J. F., Estrada, K., Peters, M. J., Herrera, L., Kruithof, C. J., Duijts, L., Hofman, A., van Duijn, C. M., Uitterlinden, A. G., Jaddoe, V. W. V., & Rivadeneira, F. (2015). Challenges in conducting genome-wide association studies in highly admixed multi-ethnic populations: The Generation R Study. European Journal of Epidemiology, 30(4), 317–330. 10.1007/s10654-015-9998-4

Molenaar, P. C., Boomsma, D. I., & Dolan, C. V. (1993). A third source of developmental differences. Behavior Genetics, 23(6), 519–524. 10.1007/BF01068142

Mulder, R. H., Neumann, A., Cecil, C. A. M., Walton, E., Houtepen, L. C., Simpkin, A. J., Rijlaarsdam, J., Heijmans, B. T., Gaunt, T. R., Felix, J. F., Jaddoe, V. W. V., Bakermans-Kranenburg, M. J., Tiemeier, H., Relton, C. L., van IJzendoorn, M. H., & Suderman, M. (2021). Epigenome-wide change and variation in DNA methylation in childhood: Trajectories from birth to late adolescence. Human Molecular Genetics, 30(1), 119–134. 10.1093/hmg/ddaa280

Muthén, L. K., & Muthén, B. O. (2017). Mplus User’s Guide (Eighth). Muthén & Muthén.

Neale, M. C., & Maes, H. H. M. (2004). Methodology for Genetic Studies of Twins and Families. Kluwer Academic Publishers.

Okbay, A., Wu, Y., Wang, N., Jayashankar, H., Bennett, M., Nehzati, S. M., Sidorenko, J., Kweon, H., Goldman, G., Gjorgjieva, T., Jiang, Y., Hicks, B., Tian, C., Hinds, D. A., Ahlskog, R., Magnusson, P. K. E., Oskarsson, S., Hayward, C., Campbell, A.,…Young, A. I. (2022). Polygenic prediction of educational attainment within and between families from genome-wide association analyses in 3 million individuals. Nature Genetics, 54(4), Article 4. 10.1038/s41588-022-01016-z

Plomin, R. (2024). Nonshared environment: Real but random. JCPP Advances, e12229. 10.1002/jcv2.12229

Privé, F., Arbel, J., & Vilhjálmsson, B. J. (2021). LDpred2: Better, faster, stronger. Bioinformatics, 36(22–23), 5424–5431. 10.1093/bioinformatics/btaa1029

R Core Team. (2024). R: A language and environment for statistical computing. [Computer software]. R Foundation for Statistical Computing. https://www.R-project.org/

Raffington, L., Tanksley, P. T., Sabhlok, A., Vinnik, L., Mallard, T., King, L. S., Goosby, B., Harden, K. P., & Tucker-Drob, E. M. (2023). Socially Stratified Epigenetic Profiles Are Associated With Cognitive Functioning in Children and Adolescents. Psychological Science, 34(2), 170–185. 10.1177/09567976221122760

Rohm, T., Andreas, A., Deppe, M., Eichhorn, H., Instinske, J., Klatzka, C. H., Kottwitz, A., Krell, K., Mönkediek, B., Paulus, L., Piesch, S., Ruks, M., Starr, A., Weigel, L., Diewald, M., Kandler, C., Riemann, R., & Spinath, F. M. (2023). Data from the German TwinLife Study: Genetic and Social Origins of Educational Predictors, Processes, and Outcomes. Journal of Open Psychology Data, 11(1). 10.5334/jopd.78

Saudino, K. J., Dale, P. S., Oliver, B., Petrill, S. A., Richardson, V., Rutter, M., Simonoff, E., Stevenson, J., & Plomin, R. (1998). The validity of parent-based assessment of the cognitive abilities of 2-year-olds. British Journal of Developmental Psychology, 16(3), 349–362. 10.1111/j.2044-835X.1998.tb00757.x

Selzam, S., Krapohl, E., von Stumm, S., O’Reilly, P. F., Rimfeld, K., Kovas, Y., Dale, P. S., Lee, J. J., & Plomin, R. (2017). Predicting educational achievement from DNA. Molecular Psychiatry, 22(2), 267–272. 10.1038/mp.2016.107

Selzam, S., Ritchie, S. J., Pingault, J.-B., Reynolds, C. A., O’Reilly, P. F., & Plomin, R. (2019). Comparing Within- and Between-Family Polygenic Score Prediction. American Journal of Human Genetics, 105(2), 351–363. 10.1016/j.ajhg.2019.06.006

Shenhar, B., Pridham, G., De Oliveira, T. L., Raz, N., Yang, Y., Deelen, J., Hägg, S., & Alon, U. (2026). Heritability of intrinsic human life span is about 50% when confounding factors are addressed. Science, 391(6784), 504–510. 10.1126/science.adz1187

Smith, Z. D., & Meissner, A. (2013). DNA methylation: Roles in mammalian development. Nature Reviews Genetics, 14(3), 204–220. 10.1038/nrg3354

Starr, A., Ruks, M., Giannelis, A., Willoughby, E., Pettersson, O., Disselkamp, C. K. L., Maj, C., Andreas, A., Ahlskog, R., Arseneault, L., Fisher, H. L., Forstner, A. J., Kandler, C., McGue, M., Nöthen, M. M., Oskarsson, S., Spinath, F. M., Vrieze, S., Wertz, J., & von Stumm, S. (2025). Within- and between-family genetic effects on educational achievement vary across countries and ages. Molecular Psychiatry. 10.1038/s41380-025-03342-0

Tellegen, P. J., Winkel, M., Wijnberg-Williams, B. J., & Laros, J. A. (2007). SON-R 2½-7: Snijders–Oomen Non-verbal Intelligence Test. Hogrefe.

Texas Education Agency. (2019). 2018–2019 Spring STAAR aggregate data. https://tea.texas.gov/texas-schools/accountability/data-reports-and-data-files/staar-aggregate-data

Tucker-Drob, E. M. (2019). Cognitive Aging and Dementia: A Life-Span Perspective. Annual Review of Developmental Psychology, 1(Volume 1, 2019), 177–196. 10.1146/annurev-devpsych-121318-085204

Tucker-Drob, E. M., & Briley, D. A. (2014). Continuity of genetic and environmental influences on cognition across the life span: A meta-analysis of longitudinal twin and adoption studies. Psychological Bulletin, 140(4), 949–979. 10.1037/a0035893

Tucker-Drob, E. M., Briley, D. A., & Harden, K. P. (2013). Genetic and Environmental Influences on Cognition Across Development and Context. Current Directions in Psychological Science, 22(5), 349–355. 10.1177/0963721413485087

Turley, P., Meyer, M. N., Wang, N., Cesarini, D., Hammonds, E., Martin, A. R., Neale, B. M., Rehm, H. L., Wilkins-Haug, L., Benjamin, D. J., Hyman, S., Laibson, D., & Visscher, P. M. (2021). Problems with Using Polygenic Scores to Select Embryos. The New England Journal of Medicine, 385(1), 78–86. 10.1056/NEJMsr2105065

Villicaña, S., & Bell, J. T. (2021). Genetic impacts on DNA methylation: Research findings and future perspectives. Genome Biology, 22(1), 127. 10.1186/s13059-021-02347-6

Wechsler, D. (2003). Wechsler Intelligence Scale for Children—Fourth Edition (WISC-IV).

Wechsler, D. (2011). WASI-II: Wechsler Abbreviated Scale of Intelligence. PsychCorp. https://psycnet.apa.org/doiLanding?doi=10.1037%2Ft15171-000

Wechsler, D. (2014). WISC-V: Administration and Scoring Manual.

Weiß, R. H. (2006). CFT 20-R. Grundintelligenztestskala 2—Revision. Hogrefe.

Weiß, R. H., & Osterland, J. (2012). Grundintelligenztest Skala 1—Revision: CFT 1-R. Hogrefe.

Woodcock, R. W., Mather, N., McGrew, K. S., & Wendling, B. J. (2001). Woodcock-Johnson III tests of cognitive abilities.

Zarandooz, S., & Raffington, L. (2025). Applying blood-derived epigenetic algorithms to saliva: Cross-tissue similarity of DNA-methylation indices of aging, physiology, and cognition. Clinical Epigenetics, 17(1), 61. 10.1186/s13148-025-01868-2

